# Characterisation of coastal reef fish assemblages across an estuarine-urbanisation gradient using underwater visual survey and environmental DNA metabarcoding

**DOI:** 10.1101/2024.08.27.607512

**Authors:** Yu-De Pei, Joseph Heard, Wenqian Xu, Shara K. K. Leung, Charmaine C. M. Yung, Alex S. J. Wyatt

**Author notes:** Yu-De Pei and Joseph Heard contributed equally to this study.

## Abstract

The ongoing urbanisation of coastlines around the globe jeopardises biodiversity, including coastal marine fishes. In many places, baseline data on fish communities are inadequate for understanding the ecological and conservation impacts of this urbanisation. Here, we document spatiotemporal patterns in fish diversity (at genus level) across an estuarine gradient around Hong Kong, a highly urbanised coastal megacity bordering the estuary of the Pearl River, the second largest river in the People’s Republic of China. We combined underwater visual census (UVC) with eDNA metabarcoding (eDNA) to overcome challenges associated with the high turbidity of Hong Kong’s aquatic environment that limits the capacity for visual observations. Similar to previous studies, UVC and eDNA captured different components of the fish community, sharing only 36.5 % fish genera in common. Nevertheless, we recorded 17 % of the known fish diversity and provided a comprehensive picture of patterns in fish diversity across the gradient, despite limited sampling effort. Fish richness was reduced by 1.6-to 3-fold under the highly turbid estuarine conditions found around Lantau compared to other regions. However, overall, there were only moderate changes in the fish community regionally. Seasonal variations in fish richness and assemblage structure were observed using both approaches, taken to reflect changes in fish behaviour, physiology, and naturally occurring events (i.e., spawning and recruitment) between seasons. A notable, consistent reduction in eDNA richness in the semi-enclosed Port Shelter might reflect limited exchange of water and genetic materials. A total of eleven species that had not been previously reported from Hong Kong were detected. These potentially novel species, as well as other ecologically and economically important species in Hong Kong, might be insufficiently protected from unregulated fishing activities due to the limited spatial coverage of marine protected areas.

## 1. Introduction

At the boundary between land and sea, the continuum of coastal waters holds some of the most biologically diverse, productive, and connected marine ecosystems, such as estuaries, seagrass beds, mangroves, and coral communities that provide critical foraging, breeding, and nursery habitats for coastal fishes (Hose et al. 1989; Parrish 1989; Nagelkerken 2007; Veneranta et al. 2013; Seitz et al. 2014; Sundblad and Bergström 2014; Sheaves et al. 2015; Lefcheck et al. 2019; Arceo-Carranza et al. 2021). Yet, the biodiversity of these ecosystems has been declining worldwide at an alarming rate due to accelerating coastal urbanisation in recent decades (Hughes 1994; Beck et al. 2001; Jackson et al. 2001; Lotze et al. 2006; Hughes et al. 2013; Ceballos et al. 2024). Coastal development, eutrophication, and pollution have damaged the habitat of coastal fishes, while over-exploitation of economically important fishes have led to shifts in fish community structure (Hughes 1994; Pauly et al. 1998; Jackson et al. 2001; Dulvy et al. 2003; Pandolfi et al. 2003; Cheung and Sadovy 2004; Lotze et al. 2006; Halpern et al. 2007, 2008; Hughes et al. 2013; Sundblad and Bergström 2014; Todd et al. 2019). The worlds’ coastal areas continue to be urbanised, however, we have yet to fully understand the ecological and conservational consequences for coastal fishes. Continuous and systematic monitoring programs are essential to reveal changes in coastal fish biodiversity across space and time concomitant with the progression of coastal urbanisation, but such comprehensive assessments are rarely conducted (Hughes et al. 2013; Duprey et al. 2017).

Hong Kong (22°9′14′′–22°33′44′′ N, 113°50’7′′–114°9′41′′ E) is one of the most urbanized metropolises on Earth. It is located on the eastern edge of Pearl River (PR) estuary, which flows into the northern South China Sea (SCS) through Hong Kong’s western region (Figure 1a). The territory has ∼1,189 km of coastline, occupies ∼1,651 km^2^ of water, entails ∼260 islands, and supports ∼7.5 million human inhabitants (Williams et al. 2019). Located in a subtropical climatic zone, Hong Kong is home to ∼1311 marine fish species of both temperate and tropical distribution (To and Shea 2016; Ng et al. 2017; Shea and To 2018; Yiu et al. 2022; Astudillo et al. 2023; Chung et al. 2023), which accounts for ∼60 % of the marine fish species recorded in the People’s Republic of China (Froese and Pauly 2024). Such high fish diversity is believed to be supported by a suite of connected marine habitats (e.g., coral communities, rocky reefs, and brackish waters), resulting from the high spatial and seasonal variability in hydrodynamics influenced by the PR (Morton and Wu 1975; Morton 1994; Ni and Kwok 1999; Sadovy and Cornish 2000; Cheung and Sadovy 2004; Lai et al. 2016; Yeung et al. 2021). Generally, Hong Kong’s aquatic environment transitions from more estuarine and more anthropogenically impacted in the west and gradually becomes more oceanic and less anthropogenically impacted towards the east. Such longitudinal transition in aquatic environment is primarily controlled by the seasonal variations in the spread of PR freshwater plume and coastal oceanic current intrusions between the summer wet season, usually between May and October, and winter dry season, usually between November and April (Chau 1958; Morton and Wu 1975). As a result, a transition in marine habitats from the west to east is evident, where coral cover and diversity increase with decreasing estuarine and urbanisation impacts from the PR (Morton 1994; McCorry 2002; Duprey et al. 2016; Yeung et al. 2021).

**Figure 1.**
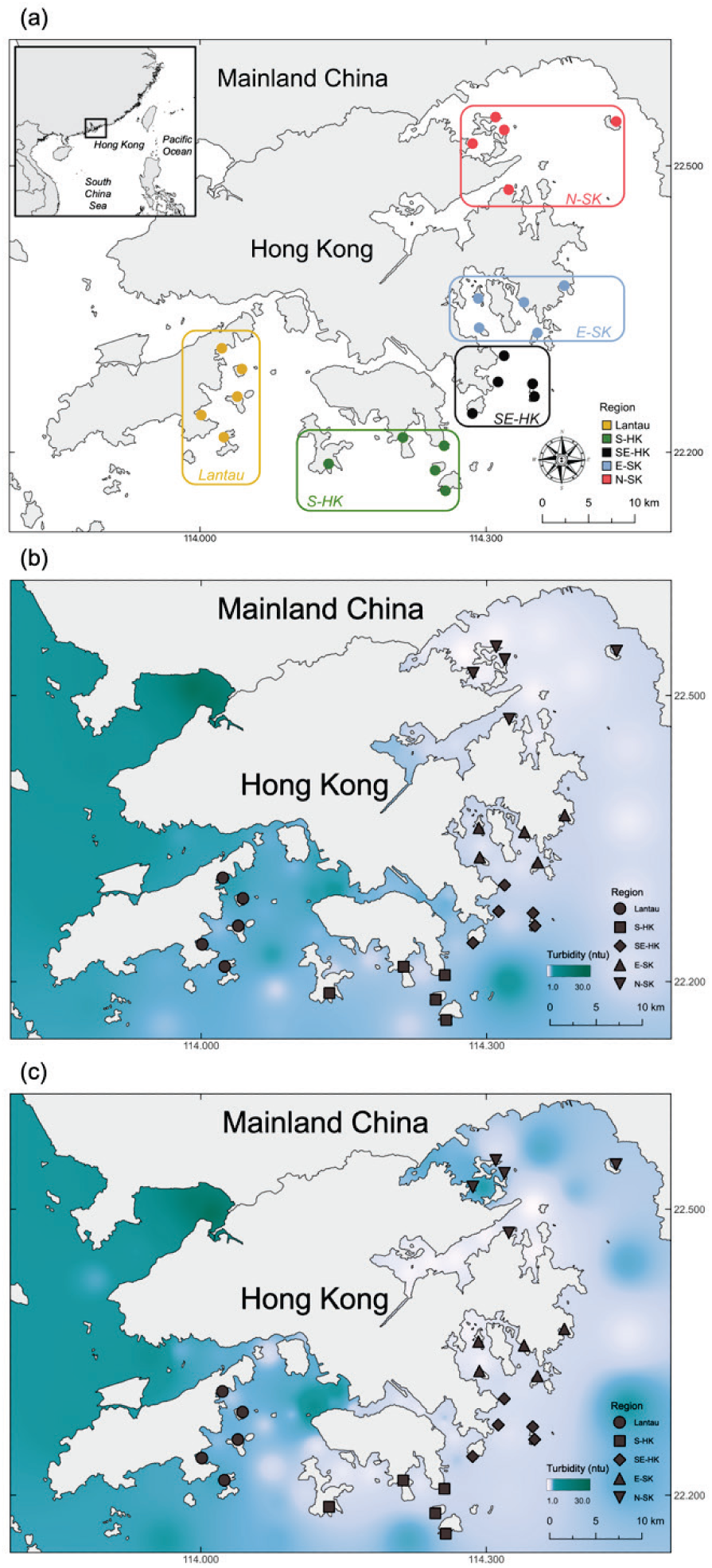
Maps showing (a) sampling sites and regions, as well as the turbidity gradients around Hong Kong waters during the (b) wet season between May and September and (c) dry season between October and April. Monthly turbidity data were pooled from 76 EPD marine water quality stations between 2017-2021 and averaged for each season. Values were then linearly interpolated over the entire area of the maps from the water quality stations using a built-in Inverse Distance Weighting (IDW) Interpolation function and represented using *singleband pseudocolor*. S-HK: South Hong Kong, SE-HK: Southeast Hong Kong, E-SK: East Sai Kung, N-SK: North Sai Kung. Maps were created using QGIS (v. 3.26.0).

The spatial and seasonal shifts in habitat and estuarine gradient are expected to affect fish community structure (Bellwood and Hughes 2001; Fabricius et al. 2005; Mallela et al. 2007; Spalding et al. 2007; Messmer et al. 2011; Neves et al. 2016; Lin et al. 2022). However, a few trawling-based studies conducted over soft bottom habitats reported contrasting and inconclusive spatial fish diversity (species richness) patterns along the region’s west-to-east transition (Leung 1997a, 1997b; Mak et al. 2021). Comparing seasons, fish diversity is generally higher during summer compared to winter, which is postulated to be a result of seasonal variations in fish behaviours, reproductive periods, as well as food and habitat availability (Sadovy and Cornish 2000; Tam and Ang 2009; Nip and Wong 2010). Despite new record species continuously being made in recent years (To and Shea 2016; Shea and To 2018; Chung et al. 2023), Hong Kong’s rich coastal fish diversity remains at risk due to continuing urbanisation (Morton 1989, 1994; Cheung and Sadovy 2004; Duprey et al. 2016, 2017, 2020), unregulated fishing activities aside from trawling and dynamite fishing, which are restricted, and incomplete closure of all fishing activities in local marine protected areas (MPAs). MPAs are also limited in size, covering only 3.7 % of Hong Kong waters (Lai et al. 2016; Chung et al. 2023). Local conservation and management efforts could therefore be greatly hampered should well-characterised baselines, particularly over coral communities and rock reef habitats, remain unavailable (Ni and Kwok 1999; Chung et al. 2023).

To quantify coastal reef fish diversity, studies generally employ a variation of catch surveys (e.g., using nets and anaesthetics), underwater visual and video surveys, or a combination of these approaches (Samoilys and Carlos 2000; Baker et al. 2016; Caldwell et al. 2016; Heard et al. 2021; Henseler and Oesterwind 2023). However, these approaches are often laborious and time consuming, and thus less economically viable or applicable to large geographical and temporal scales. Perhaps most importantly, they can be invasive and lethal. The continuing development of environmental DNA (eDNA) approaches promises to alleviate some of these limitations and has become increasingly popular to understand patterns in marine fish biodiversity (Thomsen et al. 2012; Thomsen and Willerslev 2015; Yamamoto et al. 2017; Sato et al. 2018; Seymour 2019; Miya 2022). Taxonomic information can be verified by matching gene information extracted from trace amounts of genetic materials in environmental samples to that retrieved from a curated database, which contains information of reference genetic sequences or barcodes (Miya et al. 2015; Sato et al. 2018). Earlier studies suggested that eDNA on its own will provide a more comprehensive picture of spatiotemporal variations in coastal fish diversity (Thomsen et al. 2012; Boussarie et al. 2018; Günther et al. 2018; Stat et al. 2019). However, increasing evidence has suggested that eDNA and other conventional approaches might detect different components of overall fish biodiversity, as well as yielding distinct spatiotemporal patterns in community structure (McElroy et al. 2020; West et al. 2020; Zou et al. 2020; Oka et al. 2021; Polanco Fernández et al. 2021; Valdivia-Carrillo et al. 2021; Zamani et al. 2022; Zhu et al. 2023). Multiple studies have shown that while underwater visual surveys tend to be more biased towards reef-associated fishes, eDNA appears to be able to uncover more cryptic and timid fishes and transient pelagic or demersal fishes that occur beyond reefs (Polanco Fernández et al. 2021; Valdivia-Carrillo et al. 2021; Hsu et al. 2023). The discrepancies in fish community structure across space and time, driven by the choice of approach, may in fact enable a more comprehensive characterisation of patterns in fish diversity. Nevertheless, combined visual and eDNA approaches have seldom been simultaneously evaluated, especially in locations that span strong, small-scale (∼50 km from west to east) environmental gradients, such as the estuarine-urbanisation gradient around Hong Kong.

The present study aimed to investigate the marine coastal fish diversity across Hong Kong’s estuarine-urbanisation gradient around Hong Kong between wet and dry seasons using simultaneous underwater visual census (UVC) and eDNA. We aimed to describe the local (*α*-), regional (*β*-), and territory-wide (*γ*-) diversity, as outlined by Whittaker (1972), of coastal marine fishes obtained using the two approaches and explore the seasonal patterns in the fish community across five hydrographic regions. By combining the two approaches we aimed to provide a more comprehensive picture of fish diversity at regional (site) scales, allowing us to provide guidance for fish resource management and conservation across strong environmental gradients in a highly urbanised region.

## 2. Methods

### 2.1. Study sites and turbidity gradients

Based on the hydrographic patterns, spatial benthic structure, and marine faunal distribution outlined in Morton (1989), we selected a total of 25 sites across Hong Kong from the estuarine zone in the west (eastern Lantau waters), across the middle transitional zone (southern and southeastern area of Hong Kong Island), to the eastern oceanic zone (Port Shelter and Mir’s Bay; Figure 1a). We conduced paired UVC and sampling of surface seawater for eDNA at each of the 25 sites during daytime (between 10:00 and 15:00) in May/June 2021 (wet season) and November/December 2021 (dry season).

To visualize the turbidity gradient around Hong Kong during both the wet and dry seasons, two false colour maps were created in QGIS (v. 3.26.0) based on monthly turbidity data extracted from 76 marine water quality monitoring stations of The Hong Kong Environmental Protection Department (EPD) between 2017-2021 (EPD, 2024). The turbidity data were divided into the wet (May-September) and dry (October-April) seasons.

### 2.2. Approach 1: Underwater visual census (UVC)

UVCs were conducted using a roving diver method to visually survey fish diversity (presence or absence), adopted for its high efficiency in covering a broader spatial area within a shorter period of time (Jones and Thompson 1978; Kimmel 1985; Caldwell et al. 2016; Rassweiler et al. 2020). A diver experienced in fish taxonomic identification (JH) swam freely 1 m above the benthos at a constant speed for a total of 30 minutes to record all fish species encountered on and, when visibility permitted, 5 m above the substrate (Rassweiler et al. 2020). The diver swam in a single direction along the shoreline for 10 minutes, moved 10 m further away from the shore, and continued in the opposite direction for another 10 minutes. The back-and-forth roving UVCs were completed when the 30-minute survey was completed. This ensured there was no revisiting of the surveyed area and maximised the spatial span covered. A closed-circuit rebreather was used for all underwater surveys to minimise fishes’ tendency to avoid the sound of bubbles produced by open-circuit SCUBA divers (Radford et al. 2005) and potential underestimation of fish diversity resulting from behavioural responses to heavy fishing pressure in Hong Kong (Lindfield et al. 2014; Gray et al. 2016). All surveys were conduct over coral-occupied habitats, which usually occur between 2-8 m water depth (Morton 1994; Hodgson and Yau 1997). All fish species were identified to species level, or to genus or family level if precise identification was not possible due to poor visibility.

### 2.3. Approach 2: Environmental DNA (eDNA) metabarcoding

#### 2.3.1. Sample collection and filtration

The protocol for sample collection and filtration, and downstream DNA extraction and purification, were adopted from Miya et al. (2016) with minor adjustment for the field conditions encountered in the present study. Briefly, 1 L of surface water was collected in triplicate using a pre-cleaned pitcher from the boat at each site soon after completion of the UVC. The water samples were then passed immediately through a 0.45 μm Sterivex HV pressure filter cartridge (Merck Millipore, Burlington, Massachusetts, U.S.A.) using a sterile 20 mL syringe. Upon completion of the filtration, the filter cartridges were filled with fresh RNAlater (Thermo Fisher Scientific, Waltham, Massachusetts, U.S.A.) to stop DNA degradation by denaturing proteases and RNases that might be present in the environment (Oka et al. 2021). Finally, the filter samples were placed on ice and transferred to a -80 °C freezer on returning to the lab and stored until extraction. Extraction blanks (negative controls) were also prepared by passing 1 L of ultrapure water through the filter cartridge in the lab, following the same method for field samples. The negative control samples were then processed alongside the field samples to monitor cross-contamination during the downstream experiments. Gloves were worn throughout water sampling and filter handling to reduce the risk of contamination.

#### 2.3.2. DNA extraction, purification, and PCR amplification

The eDNA retained on the filter was extracted and purified using a Qiagen DNeasy Tissue and Blood DNA extraction kit (Qiagen, Hilden, Germany), as described in Miya et al. (2016). The concentration of the extracted and purified DNA was quantified using NanoDrop spectrophotometers (BioChrom, Cambridge, Cambridgeshire, U.K.) and Qubit 4 fluorometer (Thermo Fisher Scientific, Waltham, Massachusetts, U.S.A.). The MiFish-U primers were used in PCR to amplify a hypervariable region of mitochondrial 12S rRNA gene (163-185 bp) to target teleost taxa (Miya et al. 2015). Specifically, the primers used were MiFish-U-forward (UF, 5’-GTC GGT AAA ACT CGT GCC AGC-3’) and MiFish-U-reverse (UR, 5’-CAT AGT GGG GTA TCT AAT CCC AGT TTG-3’). The recipe for a 25-μL PCR reaction included 1 μL of BSA (10 mg/ml), 0.5 μL of forward primer with barcode (10 μM), 0.5 μL of reverse (10 μM) primer with barcode, 12.5 μL of NEBNext ultra II Q5 Master Mix (New England Biolabs, Beverly, Massachusetts, U.S.A.), 9.5 μL of ultrapure molecular grade water, and 1 μL of DNA template (extracted eDNA). Cycling conditions were 95°C three minutes for denaturation, followed by 35 cycles of 20 seconds at 98°C, 15 seconds at 65°C, and 15 seconds at 72°C, with a final extension at 72°C for 10 minutes. Finally, sequencing of the PCR products and library preparation were performed using a Novaseq 6000 PE150 system (Illumina, San Diego, California, U.S.A.) by Novogene Bioinformatics Technology Co. Ltd. (Beijing, People’s Republic of China).

#### 2.3.3. Bioinformatics pipelines

The raw reads were trimmed by a 10-bp running window to remove low-quality sequence ends at a *Phred* quality of 25 using *Sickle* (Joshi and Fass 2011). Sequences with an exact match to both primers were retained, and the primer sequences were further trimmed using *Cutadapt* (Martin 2011). Paired-end reads were merged using *Usearch* (v. 11.0.667; Edgar 2010) by discarding pairs when alignment was shorter than 20 bp (*-fastq_minovlen* 20) and with a minimum 160 bp length for the Mifish amplicons’ merged sequence. The merged reads were filtered by removing those with maximum error rate larger than 0.005 (*-fastq_maxee_rate* 0.005) and then denoised using *UNOISE3* (Edgar 2016). The denoised sequences are referred to as “ZOTUs” or zero-radius operational taxonomic units. The curated ZOTUs were clustered into 0.985 OTUs using the *cluster_smallmem* command in *Usearch* (v.11.0.667) under identity 98.5% (*-id* 0.985). The 0.985 OTUs curated by removing the sum of reads that was lower than 0.02% across all samples. For fish taxonomic assignment, curated 0.985 OTUs were queried against a customised nucleotide database (MitoFish DataBase v.8Oct23 plus all Actinopterygii’s mitochondrion reads from NCBI v.28Oct23; Iwasaki et al. 2013) using Nucleotide Basic Local Alignment Search Tool (BLASTN) with an e-value threshold of 10^-5^ and alignment lengths longer than 160 bp. Finally, the top 10 best BLAST hits underwent a lowest common ancestor (LCA) calculation using *eDNAFlow* (Mousavi-Derazmahalleh et al. 2021) to determine their taxonomic information. Specifically, the curated 0.985 OTUs were assigned to species level when LCA results met or exceeded a 98.5% BLASTn identity threshold (*--lca_pid* 98.5) and 100% query coverage (*--lca_qcov* 100). An identified 0.985 OTU that could only be assigned to genus level, but not to species level, was named after its corresponding genus, followed by *“spp*” with a sequential number (e.g., *Thalassoma_spp.1*). Similarly, an identified 0.985 OTU that could only be assigned to family level, but not to genus level, or to order level, but not to family level, was named after its corresponding family or order, followed by *“sp.*” and *“f_sp”* with a sequential number, respectively (e.g., *Sparidae_sp.1* and *Pleuronectiformes_f_sp.1*). Under 100% query coverage *(--lca_qcov* 100), the curated 0.985 OTUs were assigned to the genus level, when LCA results met or exceeded a 97% BLASTn identity *(--lca_pid* 97); to family level when LCA results met or exceeded a 95% BLASTn identity, to order level when LCA results met or exceeded a 90% BLASTn identity, and to class level when LCA results met or exceeded 85% match (West et al. 2020). Curated 0.985 OTUs with BLASTn identity less than 80% and query coverage less than 90% were categorized as “non-fish” (Sato et al. 2018). Finally, multiple identified 0.985 OTUs assigned to the same species were combined into one. The raw sequence data were deposited in the National Centre for Biotechnology Information (NCBI) Sequence Read Archive (SRA) under the BioProject accession number PRJNA1147450 (https://www.ncbi.nlm.nih.gov/bioproject/).

### 2.4. Data Analyses

All fish records obtained from UVC and eDNA were first checked against the FishBase for latest accepted scientific names (Froese and Pauly 2024). Thereafter, the updated list of species was checked for their local records against the Hong Kong Register of Marine Species (HKRMS), a database for marine species found in Hong Kong waters (Astudillo et al. 2023), as well as To and Shea (2016), Shea and To (2018), Yiu et al. (2022), and Chung et al. (2023) that described new records of fish species in Hong Kong waters that were not yet included in the HKRMS. Three individuals recorded as “*Unidentifiable sp.*” from Hei Ling Chau (Lantau), Chi Ma Wan (Lantau), and Bluff Island (E-SK) during the UVC in the wet season were excluded from all analyses and statistics to avoid ambiguity surrounding data interpretation. All analyses were performed in R (v. 4.4.0; R Core Team 2024).

#### 2.4.1. Territory-wide diversity (γ-diversity)

The total number of fish taxa recorded from our study sites were compiled using data collected using UVC and eDNA. Venn diagrams were drawn using *VennDiagram* (v. 1.7.3; Chen and Boutros 2011) to investigate shared and unique taxa detected by the two sampling approaches at genus level. Fish taxa that could be ascertained to at least genus level were included for genus level comparison. Three levels of vertical position in the water column (e.g., benthic, benthopelagic, and pelagic) were assigned to each fish species based on information retrieved from FishBase (Froese and Pauly 2024) to help understand the preferential detectability of sampling approaches to certain groups of fishes. Finally, species rarefaction and extrapolation curves were drawn using *iNEXT* (v. 3.0.1) with Hill number = 0 and species by sampling-units incidence matrix to compare the territory’s fish species pool detectability between the two sampling approaches (Chao et al. 2014).

#### 2.4.2. Spatiotemporal patterns of fish generic richness (α-diversity)

To compare the spatiotemporal patterns in *α*-diversity captured by the two sampling approaches at genus level, the total number of fish genera was calculated for each region, sampling season, and approach. Means and standard deviations (std) were also calculated for each region for both sampling seasons combined and in each sampling season for each approach. The total number and mean ± std of fish genus were then visualised using boxplots. We ran a Kruskal-Wallis rank sum test (Kruskal-Wallis H test; Kruskal and Wallis 1952) using *kruskal.test* on fish generic richness using data pooled from the two methods with “approach”, “region”, and “season” as factors to test for differences between sampling approaches. Following a significant approach effect, the spatial differences in fish generic richness were then separately examined for data considering both sampling seasons, as well as for each season separately for each sampling approach. Either a two-way analysis of variance (two-way ANOVA; Yates 1934) using *aov* or a Kruskal-Wallis H test was run depending on whether data normality was confirmed statistically by Shapiro–Wilk test (*p* > 0.05; Shaphiro and Wilk 1965) using *shapiro.test* and visually by qqplot (Gnanadesikan and Wilk 1968) using *ggqqplot*. A *post hoc* pairwise Dunn test (Dunn 1961) using *dunn.test* or Tukey’s Honest Significant Difference test (Tukey’s HSD test; Tukey 1949) using *TukeyHSD* was run accordingly for the respective tests to examine between-group differences. Difference in fish generic richness within a region but between the two sampling seasons were statistically confirmed by Mann-Whitney U test using *wilcox.test* (Mann and Whitney 1947). Lastly, Venn diagrams were drawn for each sampling approach to examine shared and unique fishes to the five regions and to the two sampling seasons.

#### 2.4.3. Spatiotemporal patterns of fish composition (β-diversity)

Firstly, PERMANOVA (Anderson 2001) and PERMDISP (Anderson and Walsh 2013) tests based on the Jaccard dissimilarity (Jaccard 1912) of presence-absence data were run based on 9999 permutations using *adnois2* and *betadisper* with “approach”, “region”, and “season” as fixed factors to test for differences in fish compositions between sampling approaches. These were followed by a nonparametric multidimensional scaling (nMDS) created using *metaMDS* for visualising the differences between sampling approaches (Rabinowitz 1975). Jaccard dissimilarity is an unbiased measure of dissimilarity (i.e., the proportion of unique taxa to the total taxa pool) of two communities, and therefore commonly used with presence-absence data (Koleff et al. 2003). After confirming a significant approach effect on fish composition by PERMANOVA and PERMDISP, the spatiotemporal patterns of fish compositions were separately examined for each sampling approach. The statistical analyses included PERMANOVA and PERMDISP for investigating regional differences for both sampling seasons combined and separately, as well as nMDS for visualising the results. Indicator species analysis (ISA; Dufrêne and Legendre 1997) was then conducted, based on 999 permutations using *multipatt* with *r.g* function, to identify indicator fish genera that can best contribute to the differences in fish composition between regions. Finally, to better understand whether changes in the observed fish diversity across region and sampling seasons were due to differences in generic richness (nestedness), composition (replacement), or both, total beta diversity (BD_total_) and its two sub-divisions, nestedness (BD_nest_) and replacement (BD_repl_), across sites were computed based on Podani calculation of Jaccard dissimilarities of the presence-absence data with 999 permutations for each sampling approach using *beta.div.comp* (Legendre et al. 2005; Podani and Schmera 2011; Legendre and De Cáceres 2013; Legendre 2014).

## 3. Results

### 3.1. Variations in overall fish diversity – UVC and eDNA metabarcoding

#### 3.1.1. Territory-wide fish diversity (γ-diversity)

Collectively, the UVC and eDNA detected a total of 208 fish species, belonging to 115 genera, 58 families, and 28 orders, across all sites and the two seasons (Supplementary Table 2, 3). Separately, the UVC recorded a total of 117 species belonging to 73 genera and 41 families, while eDNA yielded a total of 118 species belonging to 84 genera and 41 families from a summation of 1,717,692 reads (Supplementary Table 2, 3, 4). Among the 117 species obtained using UVC, 96 were identified to species level, while 11 and 10 could only be identified to genus and family levels, respectively (Supplementary Table 2, 3, 4). Among the 118 species detected using eDNA, a total of 81 were successfully assigned to species level, 28 to genus level, and 9 to family level and above (Supplementary Table 2, 3, 4). Based on the 150 fish species that we could assign to specific species names, we captured ∼11 % of the known marine fish diversity in Hong Kong (To and Shea 2016; Ng et al. 2017; Shea and To 2018; Yiu et al. 2022; Chung et al. 2023). Due to a significant portion of fish species unable to be identified to species level in both approaches (UVC: 18 %; eDNA: 31 %), the follow-up statistical analyses proceeded at genus level (with fish identified to family level or above removed), unless otherwise mentioned.

Rarefaction curves suggested adequate sampling effort to capture fish diversity at “species level” for eDNA but not for UVC, which did not reach a plateau at 50 sites (Figure 2). The two-dimensional Venn diagram revealed that the two sampling approaches detected distinct groups of fishes (Figure 3). Of all fish genera, 36.5 % (42 genera) were detected by both sampling approaches and the rest, 63.5 % (73 genera), were unique to one approach. Benthic (26.4 % or 20 genera) and benthopelagic (72.6 % or 53 genera) fishes that are mostly reef-associated made up all fishes found using UVC. By contrast, eDNA revealed a mixture of 15.5 % of benthic (11 reef-associated and 3 demersal genera) and 61.9 % benthopelagic (38 reef-associated and 13 demersal genera) fish genera, making up 77.4 % of the total fish genera detected (Figure 3; Supplementary Table 3). The rest of the eDNA detections, 22.6 % (or 19 genera), were transient pelagic fishes not detected using UVC (Figure 3; Supplementary Table 3).

**Figure 2.**
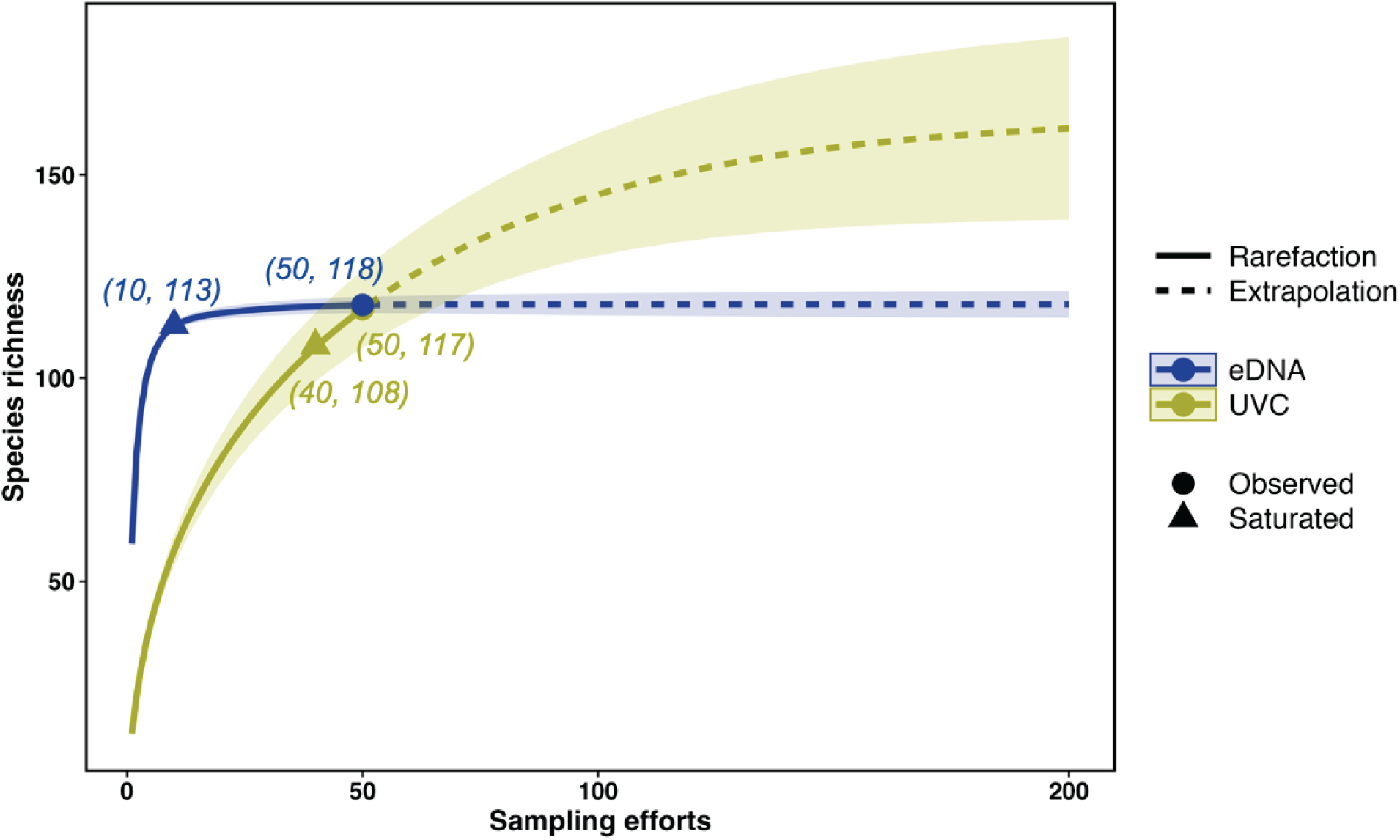
Species rarefaction curves showing species richness against sampling effort for underwater visual census (UVC in olive) and eDNA metabarcoding (eDNA in navy). Interpolated data represented by solid lines (from 0 to 50) and extrapolated data represented by dashed lines (from 51 to 200). The observed total number of species are represented as circles and the points where species detection saturated (<1% increase of species number between two consecutive sampling efforts) are represented as triangles.

**Figure 3.**
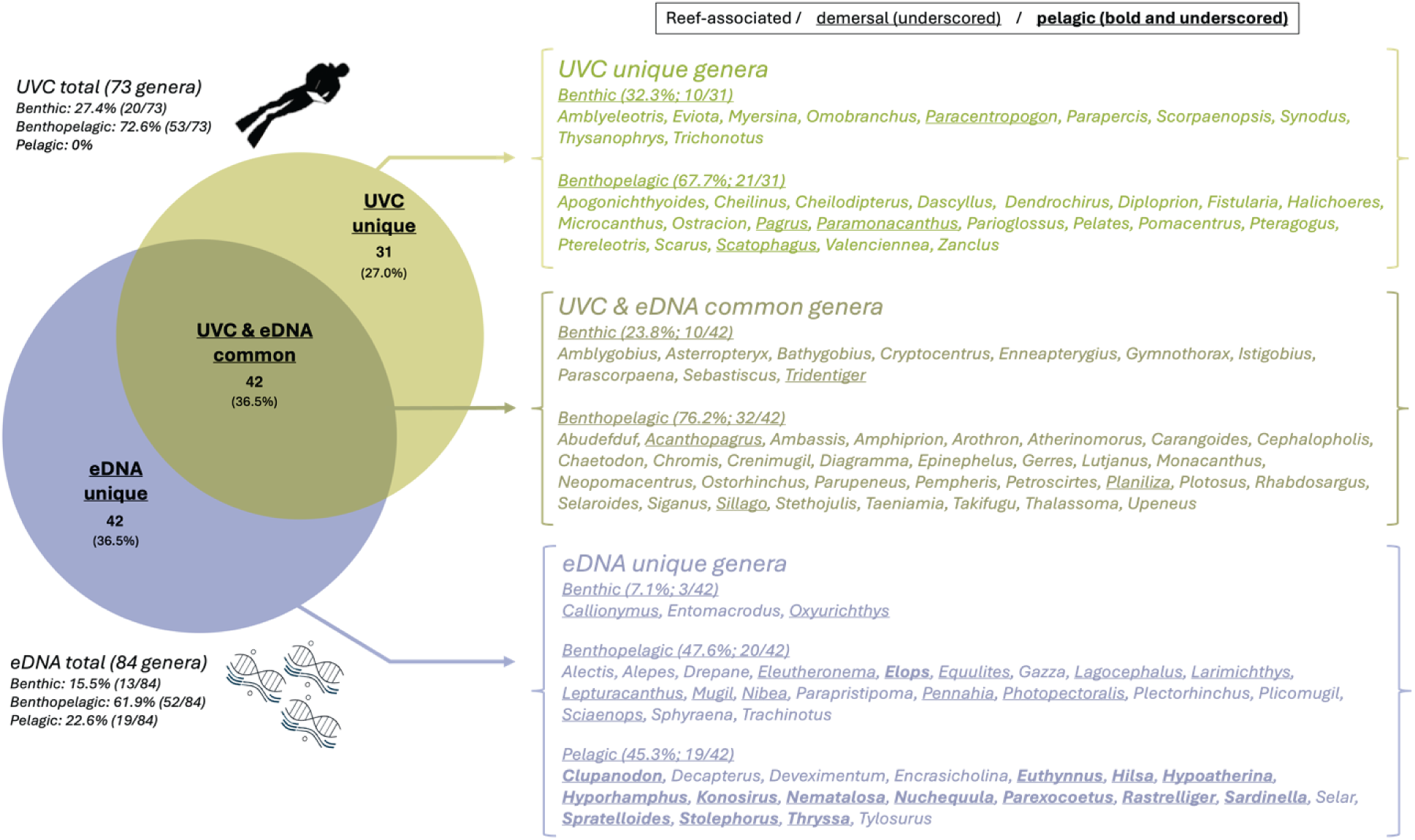
A two-dimensional Venn diagram showing the number and lists of fish genera uniquely or commonly detected by underwater visual census (UVC in olive) and eDNA metabarcoding (eDNA in navy).

#### 3.1.2. Fish generic richness (α-diversity) and assemblage structure (β-diversity)

Overall, eDNA yielded significantly higher generic richness compared to UVC (Kruskal-Wallis H test: *H* = 71.79, *df* = 1, *p* < 0.001; Supplementary Table 4). Fish assemblage structures differed significantly between approaches (PERMANOVA, *F_1.99_* = 55.13, *p* < 0.001; Supplementary Figure 1). Aside from that, fish assemblages found during UVC were also significantly more variable compared to those obtained using eDNA (PERMDISP, *p* < 0.001). The total *β*-diversity (BDtotal) across sites was higher for UVC (BD_total_ = 0.41) compared to eDNA (BD_total_ = 0.27), which is in good agreement with the PERMANOVA, PERMDISP and nMDS results. A division of total *β*-diversity in nestedness (BD_nest_) and replacement (BD_repl_) found lower values for both matrices for eDNA (BD_nest_ = 0.10; BD_repl_ = 0.17) compared to UVC (BD_nest_ = 0.21; BD_repl_ = 0.20) across regions. The ratios of BD_nest_ and BD_repl_ to the BD_total_ were roughly ∼40 % and ∼60 %, respectively for eDNA, while both parameters were ∼50 % for UVC. This implies that variations in fish compositions (replacement), rather than variations in the number of fish genera (nestedness), play a more prominent role in shaping the spatial patterns in fish assemblages evident with eDNA, but replacement and nestedness were roughly equally for UVC. Considering that the two approaches detected quite different groups of fishes (Figure 3), leading to significant variations in both fish generic richness and composition structure, we proceeded by separately examining the spatiotemporal patterns of fish generic richness and assemblage structure for each approach.

### 3.2. The spatiotemporal variations in fish diversity – UVC

#### 3.2.1. Variations in fish generic richness (α-diversity)

Fish generic richness obtained using UVC differed significantly among regions (Kruskal-Wallis H test, *H* = 21.33, *df* = 4, *p* < 0.001). The most estuarine Lantau recorded significantly lower generic richness on average compared to S-HK, E-SK, and N-SK for data combined across seasons (Dunn test, *p* < 0.05). Indeed, we reported an average of 3.8 ± 2.3 genera and a total of 19 genera for Lantau (Figure 4a; Supplementary Table 4). These figures are lower compared to an average of 10.2 ± 5.7–12.0 ± 6.2 genera and a total of 37–38 genera reported for S-HK, SE-HK, and E-SK, and are even lower when compared to an average of 16.4 ± 6.9 genera and a total of 52 genera reported for N-SK (Figure 4a; Supplementary Table 4). The notable reduction in richness found in Lantau were largely due to the lack of commonly found genera (43.8 %, 32 out of 73) between the four other regions, as well as the bulk of unique genera (30.1 %, 22 out of 73) to one of the four other regions (Supplementary Figure 2a), and 90.7 % of which (49 out of 54) associates with reef and coral habitats.

**Figure 4.**
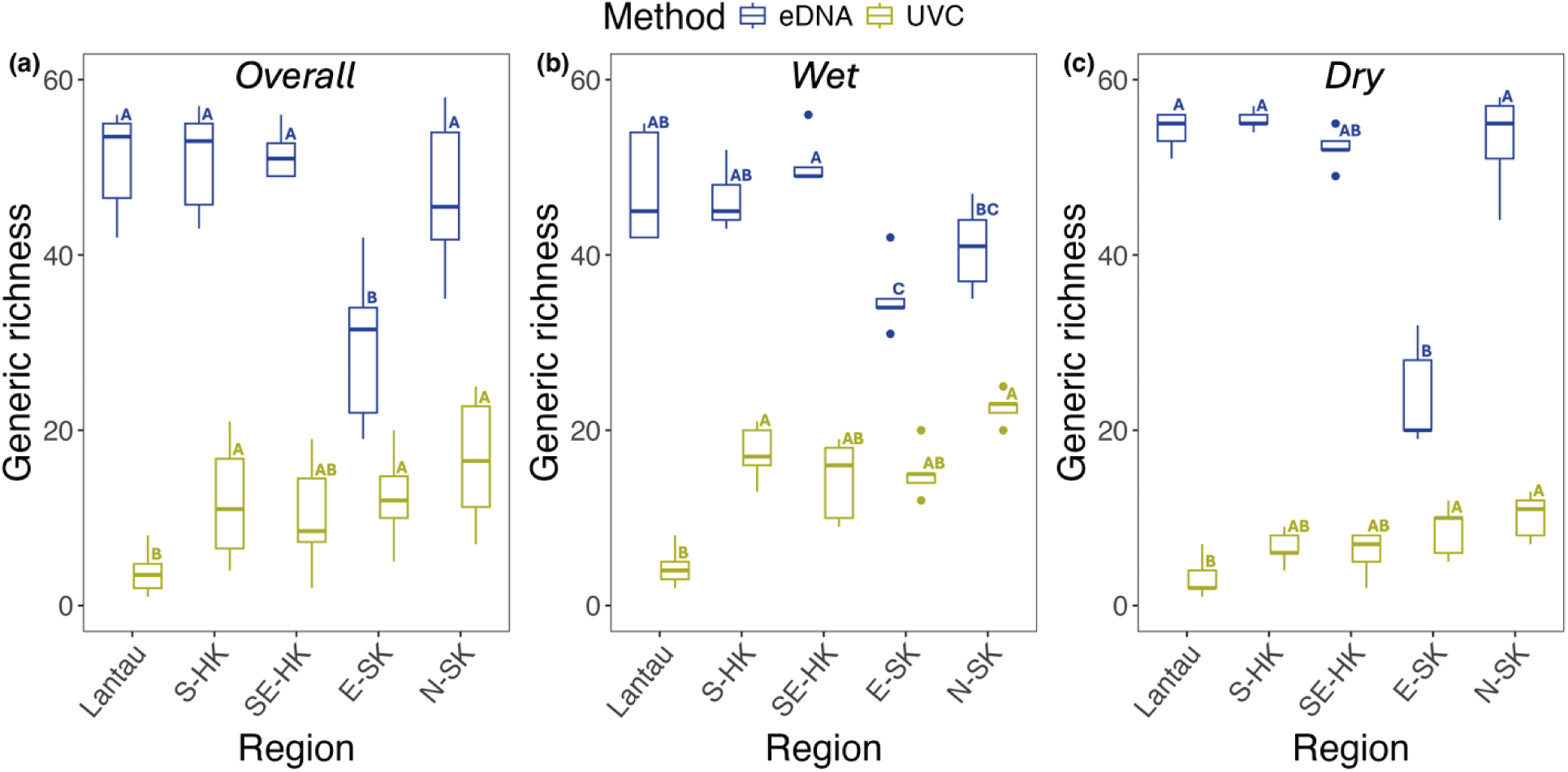
Boxplots showing differences in the fish generic richness observed between regions during both sampling seasons and separately for the wet (May/June) and dry (November/December) seasons using (a-c) underwater visual census (UVC in olive) or (d-f) eDNA metabarcoding (eDNA in navy). Boxes display median values (horizontal line) with upper and lower quartile ranges (upper and lower end of box), and maximum and minimum values (whiskers). Different lower-case letters indicate the significance groupings (at *p* < 0.05) for the mean generic richness among regions but within approach. S-HK: South Hong Kong, SE-HK: Southeast Hong Kong, E-SK: East Sai Kung, N-SK: North Sai Kung.

An overall temporal effect was also confirmed (Kruskal-Wallis H test, *H* = 16.34, *df* = 1, *p* < 0.001). Nearly twice the average number and total number of genera from each region were recorded in the wet season compared to the dry season (Mann-Whitney U test, *p* < 0.05), except for Lantau where the seasonal differences were marginal (Mann-Whitney U test, *p* > 0.05; Figure 4b, 4c; Supplementary Table 4a). This is reflected in a two-dimensional Venn diagram for seasons, where a larger number of genera were recorded as unique to the wet (30 genera) compared to dry (6 genera) season (Supplementary Figure 3a). Yet, a pool of 37 common genera between the seasons suggested the seasonal variations in fish richness might be due to both changes in fish composition (replacement) between the two seasons and more genera found during the wet season (Supplementary Figure 3a). The spatial patterns of generic richness for either the wet (one-way ANOVA, *p* < 0.001) or dry (one-way ANOVA, *p* < 0.01) season conformed relatively well with that considering both seasons above, although such spatial pattern appears to be less prominent in the dry season. In the wet season, significantly fewer fish genera were found in Lantau compared to other regions (Tukey’s HSD, *p* < 0.001), while SE-HK (Tukey’s HSD, *p* < 0.01) and E-SK (Tukey’s HSD, *p* < 0.05) each recorded significantly fewer fish genera compared to N-SK (Figure 4b). Whereas significantly fewer genera were only evident between Lantau and E-SK (Tukey’s HSD, *p* < 0.05) and N-SK (Tukey’s HSD, *p* < 0.01) in the dry season (Figure 4c).

#### 3.2.2. Variations of fish assemblage structure (β-diversity)

The compositions and variabilities of fish assemblage differed significantly between regions (PERMANOVA, *F_4.49_* = 3.19, *p* < 0.001; PERMDISP, *p* < 0.01) and seasons (PERMANOVA, *F_1,49_* = 2.89, *p* < 0.001; PERMDISP, *p* < 0.05), with also a significant interaction (PERMANOVA, *F_4,49_* = 1.31, *p* < 0.05) reported between the region and season (Supplementary Table 5a, 6a). Clear variabilities in fish assemblage structure (spread and location of the data points) can be seen between Lantau and the other regions regardless of sampling seasons, and within the same region between seasons (Figure 5a). However, differences in fish assemblages between regions became less prominent when investigated separately for the two sampling seasons (*pairwiseadonis*, *p* < 0.05), despite significant overall regional effects found for the wet (PERMANOVA, *F_4,24_* = 2.40, *p* < 0.001) and dry (PERMANOVA, *F_4,24_* = 2.13, *p* < 0.001) season (Supplementary Table 5a, 6b, 6c). Notable differences in the variability of fish composition were found for the wet season only (PERMDISP, *p* < 0.05), where fish compositions were more variable in Lantau compared to S-HK, E-SK, and N-SK (*betadisper*, *p* < 0.05; Supplementary Table 5a, 6b, 6c). This is echoed by our ISA results, which showed that some reef-associated benthic or benthopelagic fish genera displaying strong affinity to one or a few regions that do not include Lantau in mostly just the wet season (Supplementary Table 7). For instance, benthopelagic *Rhabdosargus* and *Scarus*, and benthic *Petroscirtes*, *Parioglossus* and *Ptereleotris* showed stronger affinity for N-SK in the wet season. While reef-associated benthopelagic *Apogonichthyoides* (S-HK, SE-HK, and N-SK), *Chromis* (S-HK, SE-HK, and E-SK), *Stethojulis* (S-HK, E-SK, and N-SK), *Cephalopholis*, *Parupeneus*, and benthic *Parapercis* (S-HK, SE-HK, E-SK, and N-SK) showed strong affinity to multiple regions excluding Lantau, particularly for the wet season. Compared to that, only few reef-associated benthic or benthopelagic fish genera showed strong affinity to one or a few regions across sampling seasons (Supplementary Table 7). For instance, two benthic goby genera *Asterropteryx* and *Eviota* showed stronger affinity to N-SK, another goby genus *Amblygobius* to E-SK and N-SK, and finally the benthopelagic wrasses *Halichoeres* to S-HK, SE-HK, E-SK, and N-SK. Together, these results indicate overall strong seasonal and weaker spatial effects on fish assemblage structure based on UVC.

**Figure 5.**
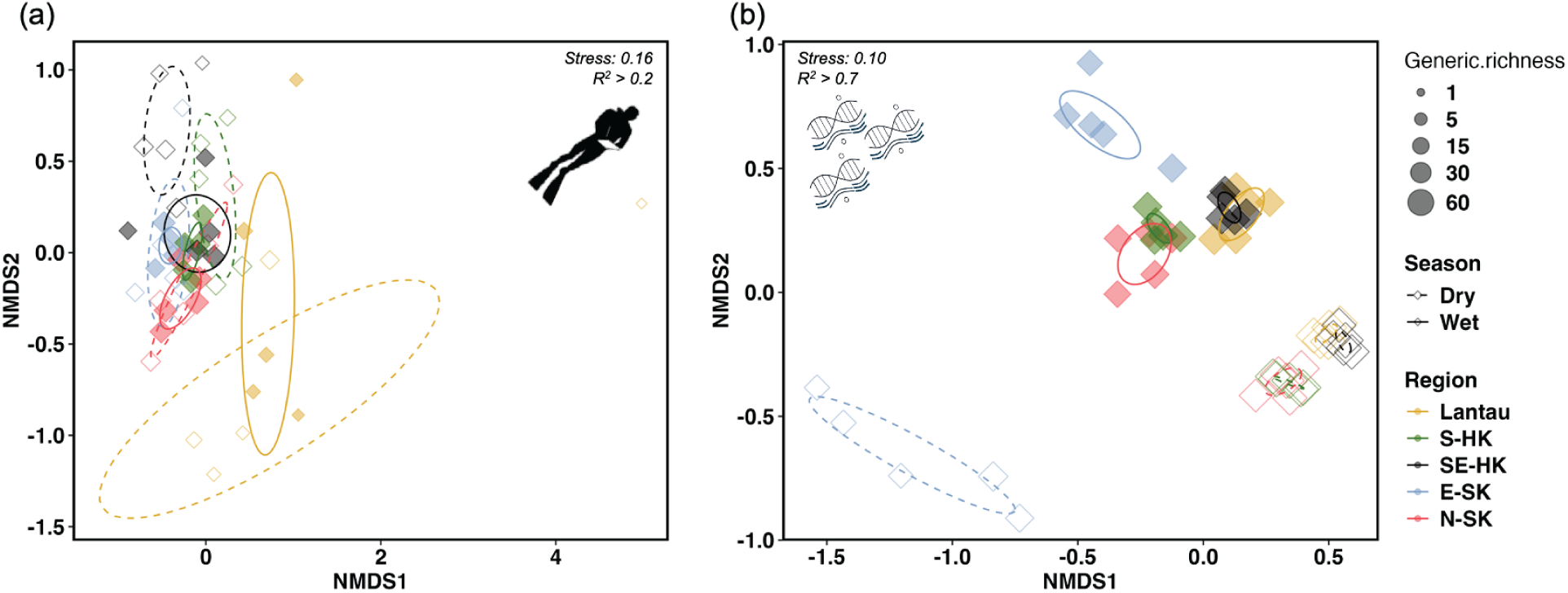
Non-metric multidimensional scaling (nMDS) plots of fish assemblages across five regions based on Jaccard’s distances of presence-absence data collected using (a) underwater visual census or (b) eDNA metabarcoding. Ellipses are drawn based on 95% confidence. The size of data points reflects fish generic richness. Wet: May/June (filled data points and solid ellipses), Dry: November/December (opened data points and dashed ellipses), Lantau (yellow), S-HK: South Hong Kong (green), SE-HK: Southeast Hong Kong (black), E-SK: East Sai Kung (sky blue), N-SK: North Sai Kung (pink).

### 3.3. The spatiotemporal variations of fish diversity – eDNA metabarcoding

#### 3.3.1. Variations of fish generic richness (α-diversity)

The fish generic richness averaged between 46.9 ± 8.2 (N-SK) and 51.4 ± 2.6 (SE-HK) and the total number of genera amounted to between 75 (Lantau) and 78 (N-SK) from our eDNA results. A pronounced spatial effect was evident (Kruskal-Wallis H test, *H* = 23.85, *df* = 4, *p* < 0.001) due to significantly lower average richness of 29.5 ± 7.7 and a reduced total of 65 genera in E-SK (Dunn test, *p* < 0.05; Figure 4a; Supplementary Table 4). Apart from E-SK, patterns in richness were relatively spatially consistent within sampling seasons. Indeed, significant spatial variation in richness can be particularly seen during the dry season (*H* = 14.92, *df* = 4, *p* < 0.01), primarily driven by reduced richness in E-SK that was > 28 genera lower than other sites on average compared to S-HK, E-SK, and N-SK during this season (Dunn test, *p* < 0.05; Figure 4c; Supplementary Table 4b). One notable exception is an insignificant difference reported between E-SK and SE-HK (Dunn test, *p* > 0.05) despite the huge difference in mean generic richness, however this is likely due to the non-parametric nature of the Kruskal-Wallis H test and Dunn test, which test for the differences in ranks among samples instead of the differences in their actual values. We, therefore, believe such difference is strong and valid. In the wet season, richness was also reduced in E-SK relative to other regions (Dunn test, *p* < 0.01), but not to N-SK (Dunn test, *p* > 0.05), although such difference is less prominent compared to that observed in the dry season (Figure 4b; Supplementary Table 4b). A reduced fish richness in E-SK can also be echoed by the absence of a total of 19 genera (12 reef-associated, 6 demersal, and 1 pelagic genus/genera) in the region that could be detected in other regions (Supplementary Figure 2b; Supplementary Table 3).

Finally, a temporal effect was also confirmed for generic richness (Kruskal-Wallis H test, *H* = 7.04, *df* = 1, *p* < 0.01). eDNA was generally able to detect 1.6 (SE-HK) to 12.2 (N-SK) more genus on average and up to 10 (N-SK) more genera in total in the dry season compared to the wet season, although significant results were not found for Lantau and SE-HK (Mann-Whitney U test, *p* > 0.05; Figure 4; Supplementary Table 4). The only exception was E-SK, where 11.7 more genus on average and 12 more genera in total were detected in the wet season (Mann-Whitney U test, *p* < 0.05; Figure 4; Supplementary Table 4). A two-dimensional Venn diagram for season showed a large pool of 53 common fish genera and the temporal variation in fish genera might be due primarily to a higher number of genera found in the dry season, which is the opposite to the seasonal pattern observed with UVC (Supplementary Figure 3b).

#### 3.3.2. Variations of fish assemblage structure (β-diversity)

Overall, significant regional (PERMANOVA, *F_4.49_* = 15.11, *p* < 0.001; PERMDISP, *p* < 0.01) and temporal (PERMANOVA, *F_1.49_* = 59.92, *p* < 0.001) effects were confirmed for the compositions and variabilities of fish assemblage, expect for an insignificant variability between sampling seasons (PERMDISP, *p* > 0.05; Supplementary Table 5b). The overall prominent compositional differences in fish assemblage were primarily driven by the distinct fish assemblage found in E-SK compared to the four other regions (*pairwiseadonis*, *p* < 0.05; Figure 5b; Supplementary Table 6a). However, such regional difference in fish compositions were not found significant when data were analysed separately for each season (*pairwiseadonis*, *p* > 0.05; Supplementary Table 5b, 6e).

The overall higher variability in fish assemblage in E-SK compared to the four other regions was supported for the dry season (*betadisper*, *p* < 0.001), but similar results were not replicated for the wet season (Supplementary Table 5b, 6f). This in turn explained the significant interaction between region and season (PERMANOVA, *F_4,49_* = 1.31, *p* < 0.05; Supplementary Table 5b).

From the ISA results, only the pelagic tenpounders *Elops* (wet season only) and reef-associated toothponies *Gazza* (dry season only) displayed strong affinity to E-SK alone, while more fish genera displayed strong affinity to multiple of other regions (Supplementary Table 7). For instance, some of the fishes that showed strong affinity across all regions except for E-SK include the benthic *Callionymus*, benthopelagic *Arothoron*, *Diagramma*, *Planiliza*, *Sillago*, *Trachinotus*, and *Upeneus*, as well as the pelagic anchovies *Thryssa* and silversides *Hypoatherina* (Supplementary Table 8). Between the two sampling seasons, we also found fish genera that generally exhibit strong affinity to either the wet (e.g. the benthopelagic *Taeniamia*, and the pelagic *Euthynnus*, *Hyporhamphus*, and *Tylosurus*) or dry (e.g., the reef-associated benthopelagic *Acanthopagrus*, *Chromis*, *Parascorpaena*, *Parapristipoma*, *Plotosus*, and *Stethojulis*, the demersal benthopelagic *Eleutheronema*, and *Lepturacanthus*, as well as the pelagic gizzard shads *Clupanodon*) seasons (Supplementary Table 8). The listed genera that showed strong affinity for the dry season above also showed weak affinity to E-SK. Finally, some fishes appeared to be well-adapted to the highly variable environmental conditions and commonly found around Hong Kong across seasons. For example, the reef-associated benthopelagic *Atherinomorus*, *Cephalopholis*, *Neopomacentrus*, and *Takifugu*, the reef-associated benthic *Enneapterygius*, *Entomacrodus*, *Sebastiscus,* as well as the benthopelagic demersal *Mugil* and pelagic *Nematalosa* showed strong affinity to all five regions (Supplementary Table 8).

## 4. Discussion

### 4.1. Overall coastal fish diversity and assemblage structures

We attempted to construct the first spatiotemporal estimates of patterns in fish diversity (*α*-, *β*-, and *γ*-diversity) across the estuarine-urbanisation gradient around Hong Kong based on genus-level data collected using UVC and eDNA. UVCs and eDNA provided similar estimates of the total numbers of fish genus overall, yet only 36.5 % of fish genera were detected by both approaches. As a result, distinct spatial patterns and close to opposing seasonal patterns in fish diversity were revealed using the two approaches. In fact, a lack of spatially distinct fish diversity patterns detected with eDNA across regions covering the environmental gradient (excluding E-SK), but not with UVC, appears to support the perspective that eDNA generally captures biodiversity at broader spatiotemporal scales (Thomsen and Willerslev 2015; Seymour 2019). Given that UVC displays higher fidelity for reef-associated fishes but can be limited by water visibility (Bozec et al. 2011), eDNA provides a complementary assessment by capturing hidden fishes, and pelagic or demersal fishes inhabiting areas beyond sight of the reef (Valdivia-Carrillo et al. 2021; Hsu et al. 2023). In marine environments, eDNA may still be detectable up to a few kilometres away from the source of release and last from a few hours to seven days (Foote et al. 2012; Thomsen et al. 2012; Baker et al. 2018). Should such findings be applicable to the condition of this study, it could partly explain the higher fish generic richness and generally (but not always) more homogenous fish assemblage structure detected with eDNA in comparison to UVC.

### 4.2. Reduced fish diversity under high-turbidity estuarine conditions and the semi-enclosed Port Shelter

Following the estuarine gradient, visible fish generic richness increased 1.6-to 2-fold in S-HK, SE-HK, and E-SK, and ∼3 fold in N-SK, compared to Lantau with a more prominent spatial pattern in the wet season compared to the dry season (Figure 4). The fish assemblage structure seen in Lantau was distinct and more variable compared to the four other regions in the wet season, although there was little evidence to suggest a drastic shift in visible fish assemblage structure among the four other regions within sampling seasons. This reduction in both generic richness and assemblage structure under estuarine conditions was not evident in eDNA data, but, as above, this could reflect the more coarse spatiotemporal resolution of eDNA as an approach. UVC might therefore be considered a higher resolution estimate that, despite methodological limitations linked to reduced water visibility, more closely reflects the localised reduction in coral-associated fish diversity at the estuarine end of Hong Kong’s environmental gradient.

Lower visible fish generic richness in the most estuarine region of Lantau can be explained by a number of environmental changes. The region’s high sedimentation rate may in turn not only affects fish behaviours and prey-predator dynamics (Wenger et al. 2013; Newport et al. 2021) but also, together with low salinity and high nutrient levels, limit coral cover, which lowers habitat and food availability (Fabricius 2005; Duprey et al. 2016, 2017; Yeung et al. 2021). Although spatial variations in visibility may also contribute to the pattern observed with UVC (Bozec et al. 2011), we were able to clearly show that certain fish genera that tend to be typically found using habitats around coral communities or rocky reefs appeared to occur less from the turbid and coral-depauperate Lantau. Identified by our ISA, these include *Apogonichthyoides*, *Parapercis*, *Parupeneus*, *Scarus*, and *Stethojulis* in the wet season, as well as *Amblygobius*, *Asterropteryx*, *Eviota*, and *Halichoeres* across both sampling seasons (Harry 2001; Mallela et al. 2007; Shibukawa and Suzuki 2007; Hernaman and Probert 2008; Randall et al. 2008; Tam and Ang 2009; Wen et al. 2010; Tornabene et al. 2015; Neves et al. 2016; To and Shea 2016). On the contrary, the small invertivorous-piscivorous scorpionfishes *Paracentropogon*, which are typically found resting on barren rocks or around soft sediments (Kwik and Lim 2020), tend to be more associated with Lantau where such habitats prevail. However, this was not statistically supported by ISA.

Some fishes appeared to be well-adapted to a range of habitats and environmental conditions despite the strong estuarine gradients that are commonly found around Hong Kong. Several of the multiple examples from the ISA include: the reef-associated planktivorous *Neopomacentrus* that has high tolerance to sedimentation compared to other damselfishes (Johansen and Jones 2013; Tarnecki et al. 2021; Esch et al. 2024), the small benthic invertivorous-piscivorous *Sebastiscus* that camouflages and ambush prey on reefs (Liu et al. 2019), the planktivorous *Atherinomorus* that usually swims in water column in a school (Kimura et al. 2001), and a few diadromous *Gerres*, *Mugil*, *Nematalosa*, *Planiliza*, *Takifugu* that traverse between freshwater and saltwater environments (Chan et al. 2024). Finally, combtooth blennies *Entomacrodus*, which have been found across a suite of shallow habitats including coral communities, tidepools, mangroves, lower reaches of rivers, oysterbeds, and even supralittoral environments throughout tropical and warm temperate waters (Hundt et al. 2014), were quite common around Hong Kong in this study. The bulk of common fishes, and a relatively small portion of fishes contributing to differences across space, may suggest that strong spatial effects along the estuarine gradient is not a universal paradigm for coastal fishes, at least for those studied here.

Finally, fish richness and assemblage structure showed a marked deviation from the expected diversity gradient based on eDNA detections in the Port Shelter (E-SK). This is rather surprising. Port Shelter is a semi-enclosed embayment known for healthy coral communities with high coral diversity and cover (Morton 1994; Duprey et al. 2016; Yeung et al. 2021), which are often positively correlated with fish diversity (Jones et al. 2004; Pratchett et al. 2011, 2018; Komyakova et al. 2013; Williamson et al. 2014; Russ et al. 2021). Although its topography makes it easier to trap nutrients and pollutants (Yung et al. 2001; Mao et al. 2011; Ho 2022; Deng et al. 2024), the water quality around the Port Shelter has substantially improved over the past two decades thanks to the continuously enhanced environmental measures, particularly the installation of sewage treatment facilities (EPD, 2024). Port Shelter also serves as traditional fishing grounds for at least nine homeports/anchorages nearby (Patchell and Cheng 2019). Yet, the fishing production (yield of fishery harvest per unit area) has dropped since a trawl ban was enforced in 2012 (Coleby and Grist 2023), with no indication of relapsing in recent years according to the latest Agriculture, Fisheries and Conservation Department’s (AFCD) Port Survey in 2021 (https://www.afcd.gov.hk; accessed August 2024). Without empirical evidence, it is currently unclear how the reef fish community responded to the relief of fishing pressure after the trawl ban in Port Shelter. However, initial recoveries of richness, abundance, or biomass has been reported for macrobenthos communities in inner and outer Port Shelter (Wang et al. 2021), as well as for macrobenthos and fish communities around Tolo Harbour, another semi-enclosed water connected to Mir’s Bay (Tao et al. 2018, 2020, 2021; Mak et al. 2021; Wang et al. 2021). If we also consider the locally high coral diversity and cover, which are themselves sensitive to poor water quality, it is unlikely that the reduced fish diversity detected in the Port Shelter in this study is a result of either suboptimal habitat / water quality or enhanced fishing pressure. The reduced flushing rates in Port Shelter may in fact hamper not just the exchange of water but also genetic materials carried in the water column from greater Mir’s Bay (Yung et al. 2001; Mao et al. 2011; Deng et al. 2024). Certainly, seasonal variations in the speed and direction of water movement greatly affect the transport of genetic materials in aquatic environments (Thomsen and Willerslev 2015; Seymour 2019). In inner Port Shelter, the current speed and flushing time can be lower than 0.15 m/s (Yung et al. 2001), and the flushing time can be ∼7-24 days in the wet season and ∼23-42 days in the dry season (Mao et al. 2011). Compared to flushing times usually less than a week in nearby open waters (Mao et al. 2011), more restricted water exchange may explain the marked reduction in fish diversity observed in Port Shelter. Moreover, a stronger northeastern monsoonal wind and reduced discharge of PR results in longer flushing times in the dry season, which may force even stronger isolation of Port Shelter from Mir’s Bay (Mao et al. 2011; Chu et al. 2022; Deng et al. 2024), thereby resulting in more reduced fish diversity and more distinct fish assemblage structure in this season. Findings in Porth Shelter therefore reinforce the importance of considering inherent constraints in eDNA detection when interpreting diversity changes across local gradients like those considered here. Some of these constraints are discussed further in Section 4.4.

### 4.3. Seasonal variations in temperature and fish behaviours affect fish diversity

The subtropical monsoonal climate around Hong Kong leads to large seasonal dynamics in the PR freshwater plume, but also to a wide range in water temperatures between seasons. Hong Kong’s sea surface temperature can drop to ∼17 °C, with the lowest temperatures measured *in situ* over coral habitats reaching as low as 13 °C to 14 °C during wintertime (ASJW, unpublished data). In the summer, temperatures rise to ∼29 °C, with occasional heatwaves reaching values of 30 °C to 32 °C (Morton 1994; Goodkin et al. 2011). The pronounces variations in water temperatures are expected to affect fish physiology, behaviour, and diversity in many ways. For instance, lower water temperatures during wintertime would decrease metabolic rates, impact physiology, and lead to reduced activity (i.e., shorter time spent and closer distance away from shelter) of certain fishes, thereby increasing mortality and making them less susceptible to observation (Fowler 1990; Mora and Ospina 2002; Feary et al. 2014; D’Agostino et al. 2020; Leriorato et al. 2021; Vaughan et al. 2021). In Hong Kong, a delayed increase in invertebrate prey availability has also been shown following the proliferation of macroalgae during wintertime, and the population of macroalgae decreases significantly in the summer until a resurgence in the next winter (Kaehler and Williams 1996; Tam and Ang 2009; Cheung-Wong et al. 2022). Such variations in food and habitat availability following the seasonal temperature variations could also affect fish occurrence and abundance (Afeworki et al. 2013; López-Pérez et al. 2013; Wilson et al. 2014; Carminatto et al. 2020; Riofrío-Lazo et al. 2022). Together, reduced activity, abundance, and occurrence will lead to lower detections with both UVC and eDNA. It is therefore not surprising that seasonality had an overall effect on fish generic richness and assemblage structure.

Although seasonality in fish diversity is expected, we found opposing patterns using UVC and eDNA. Fish generic richness was higher in the wet season and lower in the dry season for UVC, but the opposite for eDNA (except E-SK, see above). Observations similar to our UVC findings have been reported from other more localized studies in the oceanic eastern waters of Hong Kong using a variety of conventional approaches (Sadovy and Cornish 2000; Tam and Ang 2009; Cheung et al. 2023). These studies suggests that all the factors outlined above, as well as seasonal spawning and recruitment events, may contribute to increased fish diversity in the wet compared to the dry season. In this study, we visually documented a few fish genera, including the commonly found reef-associated cardinalfishes *Apogonichthyoides*, groupers *Cephalopholis*, sandperches *Parapercis*, goatfishes *Parupeneus*, and wrasses *Stethojulis*, that showed strong associations with the wet season from ISA results. Increased activity levels for foraging and movement, as well as increased abundance due to increased food availability in warmer waters, may render these fishes easier to visually observe, a possible explanation supported by our personal experience and previous studies (Wahbeh and Ajiad 1985; Sadovy and Cornish 2000; Chase et al. 2018). On the contrary, eDNA is not constrained by visual observations and can complement seasonal variations from UVC, including for certain fishes that spawn and recruit in specific seasons. For instance, the ISA results for eDNA reported a strong association of *Acanthopagrus* and *Parapristipoma* with the dry season, which aligns nicely with their empirical spawning season in the northern SCS region and Kii Peninsula, Japan, respectively (Doiuchi et al. 2007; Law and Sadovy 2017; Cheung et al. 2023; Tan et al. 2023). Certainly, seasonal variations of many other factors, such as birth rates, mortality rates, as well as ontogenetic and spawning migrations, may together with fish behaviour and spawning and recruitment timing outlined above, contribute to the fish diversity patterns. This highlights the value of complementary approaches able to detect spatiotemporal changes such as reported here for UVC and eDNA. As such approaches provide more baseline information regarding the biology and diversity of coastal reef fishes around Hong Kong and the southern coastline of China, we should be able to start verifying some of the hypotheses posed above as to what drives spatiotemporal variation in fish assemblages across the strong environmental gradient.

### 4.4. Limitations in respective approaches for characterising Hong Kong’s fish diversity

Although this study, with relatively limited effort, was able to document a reasonable proportion of the total known fish diversity for Hong Kong, some limitations need to be considered. Of the 208 species detected by both approaches, 150 could be assigned to species level, which represents ∼11 % of the known fish diversity (To and Shea 2016; Ng et al. 2017; Shea and To 2018; Yiu et al. 2022; Chung et al. 2023). Of the taxa detected with eDNA, ∼69 % (81 out of 118) could not be assigned a scientific name to species level. Of the fish species observed with UVC, ∼17 % (20 out of 117) could not be confidently assigned to species level, and aside from that, there were three unidentifiable individuals. For this reason of limited identification to species level, investigation of fish diversity patterns could only be undertaken at a relatively coarse taxonomic resolution (i.e., at genus-level). We must also consider limitations arising from poor visibility that can hamper more precise species-level observations during UVC. For instance, the record of *Crenimugil crenilabis* from UVC could conceivably belong to other species or genera under the family *Mugilidae* due to the difficulty in confidently identifying them in the field. Cryptobenthic reef fishes may contribute a rich, or even dominant, component of diversity (Brandl et al. 2018) but are less likely to be observed without the use of anaesthetic substances and thus remain relatively understudied in the region. While some of these limitations for UVC were anticipated to be addressed by eDNA, this does not appear to be the case in general. A limitation of eDNA metabarcoding is that streamlined protocols – from sample collection to bioinformatic pipelines – and reference barcoding databases suited for identifying local species found in the spatiotemporally varying Hong Kong waters are not yet available. The impact of the lack of a local reference barcode database for eDNA can be evidenced by the failed attempt to assign certain common species found in Hong Kong to species level. A few of the many examples include the retrieved *Chromis_spp.1*, *Diagramma_spp.1*, *Taeniamia_spp.1*, *Lutjanus_spp.1*, and *Labridae_sp.1*. Future studies should aim to curate a more comprehensive local reference barcode database and perhaps compare the use of different gene markers and primer design for local species of interest (Deiner et al. 2017; Stat et al. 2017; Alberdi et al. 2018; Jeunen et al. 2019). Findings from eDNA also need to acknowledge daily and seasonal changes in water flow, and seasonal variations in fish abundance, spawning timing and tissue shredding rates, as well as the longevity and vertical distribution (in the water column) of genetic materials under local conditions (Thomsen and Willerslev 2015; Seymour 2019; Miya 2022). These factors are particularly important to consider when interpreting eDNA results for coastal ecosystems with complex hydrodynamics (Nagarajan et al. 2022). In fact, as mentioned above, we suspect that the different fish community structure observed using eDNA around Port Shelter (E-SK) versus the four other regions might be partly due to a combination of these factors, particularly the limited water flow that hampers exchange of genetic material with the greater Mir’s Bay region (Yung et al. 2001; Chu et al. 2022; Deng et al. 2024). Currently, verification of the impact of these factors on fish diversity estimated using eDNA remains limited and should be a priority for robust application of eDNA metabarcoding to fish assemblage characterisation across strong environmental gradients.

### 4.5. Notes on potential novel and questionable, as well as introduced species to Hong Kong waters

Despite certain limitations that can limit taxa retrieval to species level, eDNA still holds huge potential to reveal hidden diversity, including species that might be regionally novel. From our study using eDNA, we were able to detect a total of eleven species that have not been officially reported in Hong Kong along with one introduced species (Supplementary Table 2). Among them, *Bathygobius cotticeps*, *Carangoides hedlandensis*, *Crenimugil pedaraki*, *Cryptocentrus caeruleomaculatus*, *Deveximentum indicium*, *Enneapterygius hsiojenae*, *Equulites oblongus*, *Hypoatherina woodwardi*, *Neopomacentrus anabatoides*, *Parascorpaena aurita*, *Parexocoetus mento* have been reported from waters of one or a few nearby east or southeast Asian countries, such as Korea, Japan, Taiwan, Vietnam, and the Philippines (Froese and Pauly 2024). It is therefore possible for some of these species to occur in Hong Kong, but they have not yet been documented. From personal experience (YDP), *E. hsiojenae* and *P. aurita* might have previously been encountered in Long Ke Wan in E-SK, thereby increasing the credibility of at least two of these new species records in this study (Supplementary Figure 4). The detection of *N. anabatoides* could be rather questionable, since it is reported only around the Indo-Malayan region and the Indian Ocean, and *Neopomacentrus cyanomos*, which has been frequently observed in Hong Kong (Froese and Pauly 2024), was missing from our eDNA results. We detected *Sciaenops ocellatus*, a species that is native to the northern Atlantic Ocean but universally raised in fish farms across Asia, from multiple sites of our study using eDNA (Lin et al. 2020). Considering also the multiple sighting of this species at different ontogenetic stages in Hong Kong on iNaturalist, a popular global biodiversity citizen science platforms (https://www.inaturalist.org, accessed August 2024), our findings suggests that this introduced species might have established a stable population in Hong Kong. A few more observations of potential novel species for Hong Kong were made during the UVC surveys, which recorded *Cryptocentrus cf. albidorsus*, *Cryptocentrus cf. leucostictus*, *Cryptocentrus cf. melanopus*, and *Cryptocentrus cf. sericus*, as well as *Cryptocentrus cf. caeruleomaculatus* that was also detected by eDNA. Some of these species were photographed by the primary observer of fish in this current study (Supplementary Figure 4). These are cryptobenthic fish species that have also been unofficially recorded in Hong Kong by the 114°E Reef Fish Survey (https://www.114ehkreeffish.org; accessed August 2024), while their taxonomic verification remains to be conducted. Given the poorly constructed cryptobenthic fishes diversity around the globe (Brandl et al. 2018), it is not surprising that novel species to Hong Kong or even the science community might be continuously discovered. Without substantial empirical evidence, it is not the intention of this study to report and describe novel species for Hong Kong, but our findings do point to promising directions for more in-depth investigation of these species.

### 4.6. Implications for local marine fish resource management and conservation

Morton (1976) pointed out in his seminal article, *“The problems that Hong Kong faces with regard to its various local environments are the problems of south-east Asia in microcosm”*. Certainly, the world is seeing an expansion and increasing number of coastal megacities that will attract more waves of relocation of the continuously growing human population (Small and Nicholls 2003; Barragán and De Andrés 2015). Ongoing damage to important adjacent coastal marine ecosystems seems inevitable and the establishment of coastal baselines and subsequent monitoring across urbanisation gradients are increasingly crucial to inform local and regional conversation and management plans. Linked to a huge population by the PR, Hong Kong’s marine environment is under significant urbanisation pressure and the regions’ yet to be fully documented rich biodiversity remains vulnerable (Chung et al. 2023). Hong Kong currently has a total of seven MPAs and one marine reserve administered by the AFCD but only cover 3.7 % of Hong Kong’s marine territory (https://www.afcd.gov.hk; accessed August 2024). Among them, two MPAs and the only marine reserve were established for protecting coral habitats, one for mangroves and seagrass beds, and four for important habitats of Indo-Pacific hump-backed dolphin *Sousa chinensis*, known locally as Chinese white dolphin. Except for a total of 0.01 % (0.2 km^2^) of Hong Kong’s marine territory completely protected from all fishing practices by the establishment of Cape d’Aguilar Marine Reserve, the rest of the seven MPAs still allow for traditional fishing practices by indigenous fishermen (Lai et al. 2016). The present study clearly showed strong seasonal variations and, to some extent, west-east (or other) spatial variations in fish richness and assemblage compositions. There was also evidence of differential preferences for certain seasons and habitats within individual species. As argued by Chung et al. (2023), new record species, including the yet-to-be validated records from this study, are often not found in MPAs. We further observed that some of these potential novel species display higher association to either wet (e.g., *C. hedlandensis* and *P. mento*) or dry (e.g., *B. cotticeps*, *C. caeruleomaculatus*, *C. pedaraki*, and *P. aurita*) seasons. It would not be surprising for complicated spatiotemporal heterogeneity of fish diversity to apply across a range or recorded and unrecorded ecologically and economically important, yet heavily exploited, species (Cheung and Sadovy 2004). In this regard, with the goal of 10 % territorial cover of MPAs by 2020 and 30 % by 2030 (Grorud-Colvert et al. 2021) some way off, it seems likely that the majority of local and potentially novel fish species are insufficiently protected. We would strongly advocate for re-evaluation of the appropriateness of incomplete bans on fishing activities in MPAs in respect of safeguarding Hong Kong’s already deteriorating fish diversity. Understanding how to protect the known and unknown fish diversity depends on sustained long-term monitoring of spatiotemporal patterns and fundamental understanding of species’ biology, which in turn can help predict changes in biodiversity and ecosystem functions under rapid urbanisation in the region.

### 4.7. Conclusions

In this study, we have constructed spatial and seasonal fish diversity patterns (at genus level) along an estuarine-urbanisation gradient around Hong Kong using simultaneous UVC, a conventionally used approach, and eDNA metabarcoding, an increasingly prevalent tool for diversity surveys. Although the two approaches detected different sets of fishes, together the findings provided evidence to support reduced fish richness with increasing proximity to the influence of the Pearl River estuary and intense urbanisation, yet with a relatively spatially consistent assemblage structure along the environmental gradient. This implies that despite the differences in aquatic environment and habitats that might have affected certain fishes, the bulk of common fishes are much more adaptable to changes across this gradient than perhaps assumed. Seasonality, on the contrary, was confirmed to be significant driver of differences in fish diversity (both richness and assemblage structure) in this subtropical, monsoonal environment, albeit with opposing patterns between the two approaches. We discussed how seasonal variations in fish physiology and behaviour, as well as certain naturally occurring events (spawning and recruitment) may explain the fish diversity patterns observed using either approach. The markedly reduced fish diversity around Port Shelter (E-SK) was likely a result of hydrographic isolation from the broader region, which limits the exchange of genetic materials and highlights one of the challenges of comparing eDNA results across environmental gradients. However, due to the complex and challenging environmental conditions and the lack of a genetic barcode database specifically for local fish species, we were unable to further fully constrain factors that might influence the relative performance of each approach at this time. Ongoing improvements in local barcode databases and applications of eDNA will continue to improve our understanding of fish diversity across this environmental gradient. Despite limitations, we have documented spatiotemporal patterns in fish diversity and highlighted potentially novel fish species records, which together provide an important basis for future studies and conservation and management strategies across environmental-urbanisation gradients.

## Author contributions

Conceptualisation: JH, ASJW; Data curation: JH, WX; Formal Analysis: YDP, JH, WX; Funding acquisition: ASJW; Investigation: YDP, JH, WX, SKKL; Methodology: YDP, JH, WX, ASJW; Project administration: ASJW; Resources: CY, ASJW; Software: YDP, JH, WX; Supervision: CY, ASJW; Validation: CY, ASJW; Visualisation: YDP; Writing – original draft: YDP, ASJW; Writing – review & editing: all.

## Acknowledgements

We would like to thank J. Bennett-Williams, T.B. King, A.G. Intan, and C. Skinner for assistance in water collection and filtration in the field, as well as Y. Xu for help during the molecular experiments in the lab. We are grateful for the skilled boat captains, Wai, I. Kwok, V. Tong, and M. Leung for their assistance with field logistics. YDP would also like to personally thank R.H.H. Tsang, A.Y.C. Chung, and G. Pérez-Rosales for their helpful comments during the preparation of this manuscript, as well as C.L. Lee and H. Motomura for personal communications on fish identification. Surveys were performed under permits ’(47) in AF MPD 09/3 Pt.24’, ‘(2) in AF MPD 09_3 Pt.25’, and ‘(27) in AF MPD 09_3 Pt.25’ from the Hong Kong Agriculture, Fisheries and Conservation Department. YDP and JH were supported by the Hong Kong Research Grant Council under the Hong Kong PhD Fellowship Scheme. YDP also acknowledges the support of the Bai Xian Asia Institute under the Asian Future Leaders Scholarship Program during field sampling. This study was partially supported by funding from the Hong Kong Branch of the Southern Marine Science and Engineering Laboratory Guangdong Laboratory (Guangzhou) under SMSEGL20SC01 to ASJW.

## Notes

### Competing Interest Statement

The authors have declared no competing interest.

## References

1. Afeworki Y, Videler JJ, Bruggemann JH (2013) Seasonally changing habitat use patterns among roving herbivorous fishes in the southern Red Sea: the role of temperature and algal community structure. Coral Reefs 32:475–485

2. Alberdi A, Aizpurua O, Gilbert MTP, Bohmann K (2018) Scrutinizing key steps for reliable metabarcoding of environmental samples. Methods Ecol Evol 9:134–147

3. Anderson MJ (2001) A new method for non-parametric multivariate analysis of variance. Australian Ecology 26:32–46

4. Anderson MJ, Walsh DCI (2013) PERMANOVA, ANOSIM, and the Mantel test in the face of heterogeneous dispersions: What null hypothesis are you testing? Ecol Monogr 83:557–574

5. Arceo-Carranza D, Chiappa-Carrara X, Chávez López R, Yáñez Arenas C (2021) Mangroves as feeding and breeding grounds. Mangroves: Ecology, Biodiversity and Management 63–95

6. Astudillo JC, Williams GA, Leung KMY, Cannicci S, Yasuhara M, Yau C, Qiu JW, Ang PO, To AWL, Shea SKH (2023) Hong Kong Register of Marine Species. https://www.marinespecies.org/hkrms. Accessed on 06 November 2023.

7. Baker CS, Steel D, Nieukirk S, Klinck H (2018) Environmental DNA (eDNA) from the wake of the whales: Droplet digital PCR for detection and species identification. Front Mar Sci 5:133

8. Baker DGL, Eddy TD, McIver R, Schmidt AL, Thériault M-H, Boudreau M, Courtenay SC, Lotze HK (2016) Comparative analysis of different survey methods for monitoring fish assemblages in coastal habitats. PeerJ 4:e1832

9. Barragán JM, De Andrés M (2015) Analysis and trends of the world’s coastal cities and agglomerations. Ocean Coast Manag 114:11–20

10. Beck MW, Heck KL, Able KW, Childers DL, Eggleston DB, Gillanders BM, Halpern B, Hays CG, Hoshino K, Minello TJ (2001) The identification, conservation, and management of estuarine and marine nurseries for fish and invertebrates: a better understanding of the habitats that serve as nurseries for marine species and the factors that create site-specific variability in nursery quality will improve conservation and management of these areas. Bioscience 51:633–641

11. Bellwood DR, Hughes TP (2001) Regional-Scale Assembly Rules and Biodiversity of Coral Reefs.

12. Boussarie G, Bakker J, Wangensteen OS, Mariani S, Bonnin L, Juhel J-B, Kiszka JJ, Kulbicki M, Manel S, Robbins WD (2018) Environmental DNA illuminates the dark diversity of sharks. Sci Adv 4:eaap9661

13. Bozec Y-M, Kulbicki M, Laloë F, Mou-Tham G, Gascuel D (2011) Factors affecting the detection distances of reef fish: implications for visual counts. Mar Biol 158:969–981

14. Brandl SJ, Goatley CHR, Bellwood DR, Tornabene L (2018) The hidden half: ecology and evolution of cryptobenthic fishes on coral reefs. Biological Reviews 93:1846–1873

15. Caldwell ZR, Zgliczynski BJ, Williams GJ, Sandin SA (2016) Reef fish survey techniques: assessing the potential for standardizing methodologies. PLoS One 11:e0153066

16. Carminatto AA, Rotundo MM, Butturi-Gomes D, Barrella W, Junior MP (2020) Effects of habitat complexity and temporal variation in rocky reef fish communities in the Santos estuary (SP), Brazil. Ecol Indic 108:105728

17. Ceballos G, Ehrlich PR, Barnosky AD, García A, Pringle RM, Palmer TM (2024) Accelerated modern human–induced species losses: Entering the sixth mass extinction. Sci Adv 1:e1400253

18. Chan JCF, Liew JH, Dudgeon D (2024) High spatial variability in a species-rich assemblage of diadromous fishes in Hong Kong, southern China. J Fish Biol

19. Chao A, Gotelli NJ, Hsieh TC, Sander EL, Ma KH, Colwell RK, Ellison AM (2014) Rarefaction and extrapolation with Hill numbers: a framework for sampling and estimation in species diversity studies. Ecol Monogr 84:45–67

20. Chase TJ, Nowicki JP, Coker DJ (2018) Diurnal foraging of a wild coral-reef fish Parapercis australis in relation to late-summer temperatures. J Fish Biol 93:153–158

21. Chau YK (1958) Some hydrological features of the surface water of Pearl River estuary between Hong Kong and Macau. Hong Kong University Fisheries Journal 2:37–42

22. Chen H, Boutros PC (2011) VennDiagram: a package for the generation of highly-customizable Venn and Euler diagrams in R. BMC Bioinformatics 12:1–7

23. Cheung CCY, Leung RKL, Law CSW, Cheng MCF, Ho KKY, Leung PTY, Astudillo JC, Leung KMY (2023) Juvenile fish communities in coastal soft-bottom and shallow water habitats at the Tolo Harbour and Channel in Hong Kong, South China. Reg Stud Mar Sci 62:

24. Cheung WWL, Sadovy Y (2004) Retrospective evaluation of data-limited fisheries: a case from Hong Kong. Rev Fish Biol Fish 14:181–206

25. Cheung-Wong RWY, Dytnerski JK, Gotama R, Hemraj DA, Russell BD (2022) Seasonal patterns of macroalgal and sessile invertebrate communities in a monsoonal marine ecosystem. Estuar Coast Shelf Sci 275:107962

26. Chu N, Liu G, Xu J, Yao P, Du Y, Liu Z, Cai Z (2022) Hydrodynamical transport structure and lagrangian connectivity of circulations in the Pearl River Estuary. Front Mar Sci 9:996551

27. Chung A, See GCL, Lam SY, Yiu WH, Shea SKH (2023) Thirty-one new records of reef fish species for Hong Kong waters. Journal of the Marine Biological Association of the United Kingdom 103:

28. Coleby AM, Grist EPM (2023) Fishing production and fishing changes in Hong Kong after the ground trawl ban of 31st December 2012: a geospatial evaluation. Journal of Geographical Research 6:39–53

29. D’Agostino D, Burt JA, Reader T, Vaughan GO, Chapman BB, Santinelli V, Cavalcante GH, Feary DA (2020) The influence of thermal extremes on coral reef fish behaviour in the Arabian/Persian Gulf. Coral Reefs 39:733–744

30. Deiner K, Bik HM, Mächler E, Seymour M, Lacoursière-Roussel A, Altermatt F, Creer S, Bista I, Lodge DM, De Vere N (2017) Environmental DNA metabarcoding: Transforming how we survey animal and plant communities. Mol Ecol 26:5872–5895

31. Deng T, Duan H-F, Keramat A (2024) Water exchange capability induced by seasonal and regional variability: Assessment of Hong Kong waters. Mar Pollut Bull 206:116732

32. Doiuchi R, Kokubo T, Ogawa M (2007) Age and growth of threeline grunt Parapristipoma trilineatum along the south-western coast of Kii Peninsula, Japan. Fisheries Science 73:777– 783

33. Dufrêne M, Legendre P (1997) Species assemblages and indicator species: the need for a flexible asymmetrical approach. Ecol Monogr 67:345–366

34. Dulvy NK, Sadovy Y, Reynolds JD (2003) Extinction vulnerability in marine populations. Fish and Fisheries 4:25–64

35. Dunn OJ (1961) Multiple comparisons among means. J Am Stat Assoc 56:52–64

36. Duprey NN, McIlroy SE, Ng TPT, Thompson PD, Kim T, Wong JCY, Wong CWM, Husa SM, Li SMH, Williams GA, Baker DM (2017) Facing a wicked problem with optimism: issues and priorities for coral conservation in Hong Kong. Biodivers Conserv 26:2521–2545

37. Duprey NN, Wang TX, Kim T, Cybulski JD, Vonhof HB, Crutzen PJ, Haug GH, Sigman DM, Martínez-García A, Baker DM (2020) Megacity development and the demise of coastal coral communities: Evidence from coral skeleton δ15N records in the Pearl River estuary. Glob Chang Biol 26:1338–1353

38. Duprey NN, Yasuhara M, Baker DM (2016) Reefs of tomorrow: eutrophication reduces coral biodiversity in an urbanized seascape. Glob Chang Biol 22:3550–3565

39. Edgar R (2010) Usearch.

40. Edgar RC (2016) UNOISE2: improved error-correction for Illumina 16S and ITS amplicon sequencing. BioRxiv 081257

41. Esch MM, Jarnevich CS, Simões N, McClanahan TR, Harborne AR (2024) Modeling the potential spread of the non-native regal demoiselle, Neopomacentrus cyanomos, in the western Atlantic. Coral Reefs 43:641–653

42. Fabricius K, De’ath G, McCook L, Turak E, Williams DM (2005) Changes in algal, coral and fish assemblages along water quality gradients on the inshore Great Barrier Reef. Mar Pollut Bull 51:384–398

43. Fabricius KE (2005) Effects of terrestrial runoff on the ecology of corals and coral reefs: review and synthesis. Mar Pollut Bull 50:125–146

44. Feary DA, Pratchett MS, J Emslie M, Fowler AM, Figueira WF, Luiz OJ, Nakamura Y, Booth DJ (2014) Latitudinal shifts in coral reef fishes: why some species do and others do not shift. Fish and Fisheries 15:593–615

45. Foote AD, Thomsen PF, Sveegaard S, Wahlberg M, Kielgast J, Kyhn LA, Salling AB, Galatius A, Orlando L, Gilbert MTP (2012) Investigating the potential use of environmental DNA (eDNA) for genetic monitoring of marine mammals.

46. Fowler AJ (1990) Spatial and temporal patterns of distribution and abundance of chaetodontid fishes at One Tree Reef, southern GBR. Marine ecology progress series Oldendorf 64:39–53

47. Froese R, Pauly D (2024) FishBase. World Wide Web electronic publication. www.fishbase.org. Accessed on 30 January 2024.

48. Gnanadesikan R, Wilk MB (1968) Probability plotting methods for the analysis of data. Biometrika 55:1–17

49. Goodkin NF, Switzer AD, McCorry D, DeVantier L, True JD, Hughen KA, Angeline N, Teng Yang T (2011) Coral communities of Hong Kong: Long-lived corals in a marginal reef environment. Mar Ecol Prog Ser 426:185–196

50. Gray AE, Williams ID, Stamoulis KA, Boland RC, Lino KC, Hauk BB, Leonard JC, Rooney JJ, Asher JM, Lopes Jr KH (2016) Comparison of reef fish survey data gathered by open and closed circuit SCUBA divers reveals differences in areas with higher fishing pressure. PLoS One 11:e0167724

51. Grorud-Colvert K, Sullivan-Stack J, Roberts C, Constant V, Horta e Costa B, Pike EP, Kingston N, Laffoley D, Sala E, Claudet J (2021) The MPA Guide: A framework to achieve global goals for the ocean. Science (1979) 373:eabf0861

52. Günther B, Knebelsberger T, Neumann H, Laakmann S, Martínez Arbizu P (2018) Metabarcoding of marine environmental DNA based on mitochondrial and nuclear genes. Sci Rep 8:14822

53. Halpern BS, Selkoe KA, Micheli F, Kappel C V (2007) Evaluating and ranking the vulnerability of global marine ecosystems to anthropogenic threats. Conservation biology 21:1301–1315

54. Halpern BS, Walbridge S, Selkoe KA, Kappel C V, Micheli F, d’Agrosa C, Bruno JF, Casey KS, Ebert C, Fox HE (2008) A global map of human impact on marine ecosystems. Science (1979) 319:948–952

55. Harry J (2001) Biodiversity of shallow reef fish assemblages in Western Australia using a rapid censusing technique. Rec West Aust Mus 20:247–270

56. Heard J, Tung W, Pei Y, Lin T, Lin C, Akamatsu T, Wen CKC (2021) Coastal development threatens Datan area supporting greatest fish diversity at Taoyuan Algal Reef, northwestern Taiwan. Aquat Conserv 31:590–604

57. Henseler C, Oesterwind D (2023) A comparison of fishing methods to sample coastal fish communities in temperate seagrass meadows. Mar Ecol Prog Ser 715:91–111

58. Hernaman V, Probert PK (2008) Spatial and temporal patterns of abundance of coral reef gobies (Teleostei: Gobiidae). J Fish Biol 72:1589–1606

59. Ho KC (2022) Overview of harmful algal blooms (red tides) in Hong Kong during 1975–2021. J Oceanol Limnol 40:2094–2106

60. Hodgson G, Yau EPM (1997) Physical and biological controls of coral communities in Hong Kong. 459–464

61. Hong Kong Environmental Protection Department (2024) Hong Kong Environmental Protection Department Marine Water Quality Data. https://cd.epic.epd.gov.hk/EPICRIVER/marine/. Accessed on 30 January 2024.

62. Hose JE, Cross JN, Smith SG, Diehl D (1989) Reproductive impairment in a fish inhabiting a contaminated coastal environment off Southern California. Environmental Pollution 57:139– 148

63. Hsu T-HT, Chen W-J, Denis V (2023) Navigating the scales of diversity in subtropical and coastal fish assemblages ascertained by eDNA and visual surveys. Ecol Indic 148:110044

64. Hughes TP (1994) Catastrophes, phase shifts, and large-scale degradation of a Caribbean coral reef. New Series 265:

65. Hughes TP, Huang H, Young MAL (2013) The wicked problem of China’s disappearing coral reefs. Conservation Biology 27:261–269

66. Hundt PJ, Iglésias SP, Hoey AS, Simons AM (2014) A multilocus molecular phylogeny of combtooth blennies (Percomorpha: Blennioidei: Blenniidae): multiple invasions of intertidal habitats. Mol Phylogenet Evol 70:47–56

67. Iwasaki W, Fukunaga T, Isagozawa R, Yamada K, Maeda Y, Satoh TP, Sado T, Mabuchi K, Takeshima H, Miya M (2013) MitoFish and MitoAnnotator: a mitochondrial genome database of fish with an accurate and automatic annotation pipeline. Mol Biol Evol 30:2531– 2540

68. Jaccard P (1912) The distribution of the flora in the alpine zone. New phytologist 11:37–50

69. Jackson JBC, Kirby MX, Berger WH, Bjorndal KA, Botsford LW, Bourque BJ, Bradbury RH, Cooke R, Erlandson J, Estes JA (2001) Historical overfishing and the recent collapse of coastal ecosystems. Science (1979) 293:629–637

70. Jeunen G, Knapp M, Spencer HG, Taylor HR, Lamare MD, Stat M, Bunce M, Gemmell NJ (2019) Species-level biodiversity assessment using marine environmental DNA metabarcoding requires protocol optimization and standardization. Ecol Evol 9:1323–1335

71. Johansen JL, Jones GP (2013) Sediment-induced turbidity impairs foraging performance and prey choice of planktivorous coral reef fishes. Ecological Applications 23:1504–1517

72. Jones GP, McCormick MI, Srinivasan M, Eagle J V (2004) Coral decline threatens fish biodiversity in marine reserves. Proceedings of the National Academy of Sciences 101:8251– 8253

73. Jones RS, Thompson MJ (1978) Comparison of Florida reef fish assemblages using a rapid visual technique. Bull Mar Sci 28:159–172

74. Joshi NA, Fass JN (2011) Sickle: a sliding-window, adaptive, quality-based trimming tool for FastQ files.

75. Kaehler S, Williams GA (1996) Distribution of algae on tropical rocky shores: spatial and temporal patterns of non-coralline encrusting algae in Hong Kong. Mar Biol 125:177–187

76. Kimmel JJ (1985) A new species-time method for visual assessment of fishes and its comparison with established methods. Environ Biol Fishes 12:23–32

77. Kimura S, Iwatsuki Y, Yoshino T (2001) Redescriptions of the Indo-West Pacific atherinid fishes, Atherinomorus endrachtensis (Quoy and Gaimard, 1825) and A. duodecimalis (Valenciennes in Cuvier and Valenciennes, 1835). Ichthyol Res 48:167–177

78. Koleff P, Gaston KJ, Lennon JJ (2003) Measuring beta diversity for presence–absence data. Journal of Animal Ecology 72:367–382

79. Komyakova V, Munday PL, Jones GP (2013) Relative importance of coral cover, habitat complexity and diversity in determining the structure of reef fish communities. PLoS One 8:e83178

80. Kruskal WH, Wallis WA (1952) Use of ranks in one-criterion variance analysis. J Am Stat Assoc 47:583–621

81. Kwik JTB, Lim KKP (2020) Scorpionfishes (Teleostei: Scorpaenoidei) of Singapore. Nature in Singapore 13:11–26

82. Lai RWS, Perkins MJ, Ho KKY, Astudillo JC, Yung MMN, Russell BD, Williams GA, Leung KMY (2016) Hong Kong’s marine environments: History, challenges and opportunities. Reg Stud Mar Sci 8:259–273

83. Law CSW, Sadovy Y (2017) Reproductive biology of black seabream Acanthopagrus schlegelii, threadfin porgy Evynnis cardinalis and red pargo Pagrus major in the northern South China Sea with consideration of fishery status and management needs. J Fish Biol 91:101–125

84. Lefcheck JS, Hughes BB, Johnson AJ, Pfirrmann BW, Rasher DB, Smyth AR, Williams BL, Beck MW, Orth RJ (2019) Are coastal habitats important nurseries? A meta-analysis. Conserv Lett 12:e12645

85. Legendre P (2014) Interpreting the replacement and richness difference components of beta diversity. Global Ecology and Biogeography 23:1324–1334

86. Legendre P, Borcard D, Peres-Neto PR (2005) Analyzing beta diversity: Partitioning the spatial variation of community composition data. Ecol Monogr 75:435–450

87. Legendre P, De Cáceres M (2013) Beta diversity as the variance of community data: dissimilarity coefficients and partitioning. Ecol Lett 16:951–963

88. Leriorato JC, Nakamura Y, Uy WH (2021) Cold thermal tolerance as a range-shift predictive trait: an essential link in the disparity of occurrence of tropical reef fishes in temperate waters. Mar Biol 168:93

89. Leung AWY (1997a) The epibenthic ichthyofauna of Tolo harbour and Hong Kong’s northeastern waters: A long term record of change. In: Morton B. (eds) Proceedings of the Eighth International Marine Biological Workshop: The Marine Flora and Fauna of Hong Kong and Southern China. Hong Kong University Press, Hong Kong, pp 463–487

90. Leung AWY (1997b) The impact of dredging and fishing on the benthic fish fauna of the southeastern waters of Hong Kong. In: Morton B. (eds) Proceedings of the Eighth International Marine Biological Workshop: The Marine Flora and Fauna of Hong Kong and Southern China. Hong Kong University Press, Hong Kong, pp 437–462

91. Lin B, Wang Y, Li J, Kang B, Fang L, Zheng L, Liu M (2020) First records of small juveniles of the red drum Sciaenops ocellatus (Linnaeus, 1766) in a subtropical mangrove habitat of China. Bioinvasions Rec 9:96–102

92. Lin YV, Hsiao WV, Chen WJ, Denis V (2022) Habitat change and its consequences on reef fish specialization in biogeographic transition zones. J Biogeogr 49:1549–1561

93. Lindfield SJ, Harvey ES, McIlwain JL, Halford AR (2014) Silent fish surveys: bubble-free diving highlights inaccuracies associated with SCUBA-based surveys in heavily fished areas. Methods Ecol Evol 5:1061–1069

94. Liu Q, Liu L, Song N, Wang X, Gao T (2019) Genetic structure in the marbled rockfish (Sebastiscus marmoratus) across most of the distribution in the northwestern Pacific. Journal of Applied Ichthyology 35:1249–1259

95. López-Pérez RA, Calderon-Aguilera LE, Zepeta-Vilchis RC, López Pérez Maldonado I, López Ortiz AM (2013) Species composition, habitat configuration and seasonal changes of coral reef fish assemblages in western M exico. Journal of Applied Ichthyology 29:437–448

96. Lotze HK, Lenihan HS, Bourque BJ, Bradbury RH, Cooke RG, Kay MC, Kidwell SM, Kirby MX, Peterson CH, Jackson JBC (2006) Depletion, degradation, and recovery potential of estuaries and coastal seas. Science (1979) 312:1806–1809

97. Mak YKY, Tao LSR, Ho VCM, Dudgeon D, Cheung WWL, Leung KMY (2021) Initial recovery of demersal fish communities in coastal waters of Hong Kong, South China, following a trawl ban. Rev Fish Biol Fish 31:989–1007

98. Mallela J, Roberts C, Harrod C, Goldspink CR (2007) Distributional patterns and community structure of Caribbean coral reef fishes within a river-impacted bay. J Fish Biol 70:523–537

99. Mann HB, Whitney DR (1947) On a test of whether one of two random variables is stochastically larger than the other. The annals of mathematical statistics 50–60

100. Mao J, Wong KTM, Lee JHW, Choi KW (2011) Tidal flushing time of marine fish culture zones in Hong Kong. China ocean engineering 25:625–643

101. Martin M (2011) Cutadapt removes adapter sequences from high-throughput sequencing reads. EMBnet journal 17:10–12

102. McCorry D (2002) Hong Kong scleractinian coral communities: status, threats, and proposals for management. PhD thesis, The University of Hong Kong

103. McElroy ME, Dressler TL, Titcomb GC, Wilson EA, Deiner K, Dudley TL, Eliason EJ, Evans NT, Gaines SD, Lafferty KD, Lamberti GA, Li Y, Lodge DM, Love MS, Mahon AR, Pfrender ME, Renshaw MA, Selkoe KA, Jerde CL (2020) Calibrating Environmental DNA Metabarcoding to Conventional Surveys for Measuring Fish Species Richness. Front Ecol Evol 8:

104. Messmer V, Jones GP, Munday PL, Holbrook SJ, Schmitt RJ, Brooks AJ (2011) Habitat biodiversity as a determinant of fish community structure on coral reefs. Ecology 92:2285– 2298

105. Miya M (2022) Environmental DNA metabarcoding: A novel method for biodiversity monitoring of marine fish communities. Ann Rev Mar Sci 14:161–185

106. Miya M, Minamoto T, Yamanaka H, Oka S, Sato K, Yamamoto S, Sado T, Doi H (2016) Use of a filter cartridge for filtration of water samples and extraction of environmental DNA. JoVE (Journal of Visualized Experiments) e54741

107. Miya M, Sato Y, Fukunaga T, Sado T, Poulsen JY, Sato K, Minamoto T, Yamamoto S, Yamanaka H, Araki H (2015) MiFish, a set of universal PCR primers for metabarcoding environmental DNA from fishes: detection of more than 230 subtropical marine species. R Soc Open Sci 2:150088

108. Mora C, Ospina A (2002) Experimental effect of cold, La Nina temperatures on the survival of reef fishes from Gorgona Island (eastern Pacific Ocean). Mar Biol 141:789–793

109. Morton B (1989) Pollution of the coastal waters of Hong Kong. Mar Pollut Bull 20:310–318

110. Morton B (1994) Hong Kong’s coral communities: Status, threats and management plans. Mar Pollut Bull 29:74–83

111. Morton B, Wu SS (1975) The hydrology of the coastal waters of Hong Kong. Environ Res 10:319– 347

112. Morton BS (1976) The Hong Kong sea-shore—an environment in crisis. Environ Conserv 3:243– 254

113. Mousavi-Derazmahalleh M, Stott A, Lines R, Peverley G, Nester G, Simpson T, Zawierta M, De La Pierre M, Bunce M, Christophersen CT (2021) eDNAFlow, an automated, reproducible and scalable workflow for analysis of environmental DNA sequences exploiting Nextflow and Singularity. Mol Ecol Resour 21:1697–1704

114. Nagarajan RP, Bedwell M, Holmes AE, Sanches T, Acuña S, Baerwald M, Barnes MA, Blankenship S, Connon RE, Deiner K (2022) Environmental DNA methods for ecological monitoring and biodiversity assessment in estuaries. Estuaries and Coasts 45:2254–2273

115. Nagelkerken I (2007) Are non-estuarine mangroves connected to coral reefs through fish migration? Bull Mar Sci 80:595–607

116. Neves LM, Teixeira-Neves TP, Pereira-Filho GH, Araujo FG (2016) The farther the better: effects of multiple environmental variables on reef fish assemblages along a distance gradient from river influences. PLoS One 11:e0166679

117. Newport C, Padget O, de Perera TB (2021) High turbidity levels alter coral reef fish movement in a foraging task. Sci Rep 11:5976

118. Ng TPT, Cheng MCF, Ho KKY, Lui GCS, Leung KMY, Williams GA (2017) Hong Kong’s rich marine biodiversity: the unseen wealth of South China’s megalopolis. Biodivers Conserv 26:23–36

119. Ni I-H, Kwok K-Y (1999) Marine Fish Fauna in Hong Kong Waters. Zool Stud 38:

120. Nip THM, Wong CK (2010) Juvenile fish assemblages in mangrove and non-mangrove soft-shore habitats in eastern Hong Kong. Zool Stud 49:760–778

121. Oka S, Doi H, Miyamoto K, Hanahara N, Sado T, Miya M (2021) Environmental DNA metabarcoding for biodiversity monitoring of a highly diverse tropical fish community in a coral reef lagoon: Estimation of species richness and detection of habitat segregation. Environmental DNA 3:55–69

122. Pandolfi JM, Bradbury RH, Sala E, Hughes TP, Bjorndal KA, Cooke RG, McArdle D, McClenachan L, Newman MJH, Paredes G (2003) Global trajectories of the long-term decline of coral reef ecosystems. Science (1979) 301:955–958

123. Parrish JD (1989) Fish communities of interacting shallow-water habitats in tropical oceanic regions. Marine ecology progress series Oldendorf 58:143–160

124. Patchell J, Cheng C (2019) Resilience of an inshore fishing population in Hong Kong: Paradox and potential for sustainable fishery policy. Mar Policy 99:157–169

125. Pauly D, Christensen V, Dalsgaard J, Froese R, Torres Jr F (1998) Fishing down marine food webs. Science (1979) 279:860–863

126. Podani J, Schmera D (2011) A new conceptual and methodological framework for exploring and explaining pattern in presence – absence data. Oikos 120:1625–1638

127. Polanco Fernández A, Marques V, Fopp F, Juhel JB, Borrero-Pérez GH, Cheutin MC, Dejean T, González Corredor JD, Acosta-Chaparro A, Hocdé R, Eme D, Maire E, Spescha M, Valentini A, Manel S, Mouillot D, Albouy C, Pellissier L (2021) Comparing environmental DNA metabarcoding and underwater visual census to monitor tropical reef fishes. Environmental DNA 3:142–156

128. Pratchett MS, Hoey AS, Wilson SK, Messmer V, Graham NAJ (2011) Changes in biodiversity and functioning of reef fish assemblages following coral bleaching and coral loss. Diversity (Basel) 3:424–452

129. Pratchett MS, Thompson CA, Hoey AS, Cowman PF, Wilson SK (2018) Effects of coral bleaching and coral loss on the structure and function of reef fish assemblages. Coral bleaching: Patterns, processes, causes and consequences 265–293

130. R Core Team (2024) A language and environment for statistical computing. Vienna, Austria: R Foundation for Statistical Computing; 2024.

131. Rabinowitz GB (1975) An introduction to nonmetric multidimensional scaling. Am J Pol Sci 343– 390

132. Radford CA, Jeffs AG, Tindle CT, Cole RG, Montgomery JC (2005) Bubbled waters: The noise generated by underwater breathing apparatus. Mar Freshw Behav Physiol 38:259–267

133. Randall JE, Senou H, Yoshino T (2008) Three new pinguipedid fishes of the genus Parapercis from Japan. Bulletin of the National Museum of Nature and Science, Series A 2:69–84

134. Rassweiler A, Dubel AK, Hernan G, Kushner DJ, Caselle JE, Sprague JL, Kui L, Lamy T, Lester SE, Miller RJ (2020) Roving divers surveying fish in fixed areas capture similar patterns in biogeography but different estimates of density when compared with belt transects. Front Mar Sci 7:272

135. Riofrío-Lazo M, Zetina-Rejón MJ, Vaca-Pita L, Murillo-Posada JC, Páez-Rosas D (2022) Fish diversity patterns along coastal habitats of the southeastern Galapagos archipelago and their relationship with environmental variables. Sci Rep 12:3604

136. Russ GR, Rizzari JR, Abesamis RA, Alcala AC (2021) Coral cover a stronger driver of reef fish trophic biomass than fishing. Ecological Applications 31:e02224

137. Sadovy Y, Cornish AS (2000) Reef fishes of Hong Kong. Hong Kong University Press

138. Samoilys MA, Carlos G (2000) Determining methods of underwater visual census for estimating the abundance of coral reef fishes. Environ Biol Fishes 57:289–304

139. Sato Y, Miya M, Fukunaga T, Sado T, Iwasaki W (2018) MitoFish and MiFish pipeline: a mitochondrial genome database of fish with an analysis pipeline for environmental DNA metabarcoding. Mol Biol Evol 35:1553–1555

140. Seitz RD, Wennhage H, Bergström U, Lipcius RN, Ysebaert T (2014) Ecological value of coastal habitats for commercially and ecologically important species. ICES Journal of Marine Science 71:648–665

141. Seymour M (2019) Rapid progression and future of environmental DNA research. Commun Biol 2:80

142. Shaphiro S, Wilk M (1965) An analysis of variance test for normality. Biometrika 52:591–611

143. Shea SKH, To AWL (2018) Ocean fifteen: New records of reef fish species in Hong Kong. Mar Biodivers Rec 11:1–9

144. Sheaves M, Baker R, Nagelkerken I, Connolly RM (2015) True value of estuarine and coastal nurseries for fish: incorporating complexity and dynamics. Estuaries and Coasts 38:401–414

145. Shibukawa K, Suzuki T (2007) Two new species of the cheek-spine goby genus Asterropteryx (Perciformes: Gobiidae: Gobiinae) from the western Pacific. Bulletin of the National Museum of Nature and Science, Series A Supplement 1:109–121

146. Small C, Nicholls RJ (2003) A global analysis of human settlement in coastal zones. J Coast Res 584–599

147. Spalding MD, Fox HE, Allen GR, Davidson N, Ferdaña ZA, Finlayson MAX, Halpern BS, Jorge MA, Lombana AL, Lourie SA (2007) Marine ecoregions of the world: a bioregionalization of coastal and shelf areas. Bioscience 57:573–583

148. Stat M, Huggett MJ, Bernasconi R, DiBattista JD, Berry TE, Newman SJ, Harvey ES, Bunce M (2017) Ecosystem biomonitoring with eDNA: metabarcoding across the tree of life in a tropical marine environment. Sci Rep 7:12240

149. Stat M, John J, DiBattista JD, Newman SJ, Bunce M, Harvey ES (2019) Combined use of eDNA metabarcoding and video surveillance for the assessment of fish biodiversity. Conservation Biology 33:196–205

150. Sundblad G, Bergström U (2014) Shoreline development and degradation of coastal fish reproduction habitats. Ambio 43:1020–1028

151. Tam TW, Ang JPO (2009) Structures, dynamics and stability of reef fish assemblages in non-reefal coral communities in Hong Kong, China. Aquat Conserv 19:301–321

152. Tan Z, Wu F, Rao Y, Pan C, Hou G, Huang H (2023) Spatial and temporal distribution of fish egg communities in the adjacent waters of Daya Bay nuclear power plant and their relationship with environmental factors. Front Mar Sci 10:1182213

153. Tao LSR, Lau DCP, Perkins MJ, Hui TTY, Yau JKC, Mak YKY, Lau ETC, Dudgeon D, Leung KMY (2020) Stable-isotope based trophic metrics reveal early recovery of tropical crustacean assemblages following a trawl ban. Ecol Indic 117:106610

154. Tao LSR, Lui GCS, Wong KJH, Hui TTY, Mak YKY, Sham RC, Yau JKC, Cheung WWL, Leung KMY (2021) Does a trawl ban benefit commercially important Decapoda and Stomatopoda in Hong Kong? Ecosystems 24:1157–1170

155. Tao LSR, Lui KKY, Lau ETC, Ho KKY, Mak YKY, Sadovy de Mitcheson Y, Leung KMY (2018) Trawl ban in a heavily exploited marine environment: Responses in population dynamics of four stomatopod species. Sci Rep 8:17876

156. Tarnecki JH, Garner SB, Patterson III WF (2021) Non-native regal demoiselle, Neopomacentrus cyanomos, presence, abundance, and habitat factors in the North-Central Gulf of Mexico. Biol Invasions 23:1681–1693

157. Thomsen PF, Kielgast J, Iversen LL, Møller PR, Rasmussen M, Willerslev E (2012) Detection of a diverse marine fish fauna using environmental DNA from seawater samples.

158. Thomsen PF, Willerslev E (2015) Environmental DNA–An emerging tool in conservation for monitoring past and present biodiversity. Biol Conserv 183:4–18

159. To AWL, Shea SKH (2016) New records of four reef fish species for Hong Kong. Mar Biodivers Rec 9:1–6

160. Todd PA, Heery EC, Loke LHL, Thurstan RH, Kotze DJ, Swan C (2019) Towards an urban marine ecology: characterizing the drivers, patterns and processes of marine ecosystems in coastal cities. Oikos 128:1215–1242

161. Tornabene L, Valdez S, Erdmann M, Pezold F (2015) Support for a ‘Center of Origin’in the Coral Triangle: Cryptic diversity, recent speciation, and local endemism in a diverse lineage of reef fishes (Gobiidae: Eviota). Mol Phylogenet Evol 82:200–210

162. Tukey JW (1949) Comparing individual means in the analysis of variance. Biometrics 99–114

163. Valdivia-Carrillo T, Rocha-Olivares A, Reyes-Bonilla H, Domínguez-Contreras JF, Munguia-Vega A (2021) Integrating eDNA metabarcoding and simultaneous underwater visual surveys to describe complex fish communities in a marine biodiversity hotspot. Mol Ecol Resour 21:1558–1574

164. Vaughan GO, Shiels HA, Burt JA (2021) Seasonal variation in reef fish assemblages in the environmentally extreme southern Persian/Arabian Gulf. Coral Reefs 40:405–416

165. Veneranta L, Hudd R, Vanhatalo J (2013) Reproduction areas of sea-spawning coregonids reflect the environment in shallow coastal waters. Mar Ecol Prog Ser 477:231–250

166. Wahbeh MI, Ajiad A (1985) The food and feeding habits of the goatfish, Parupeneus barberinus (Lacepede), from Aqaba, Jordan. J Fish Biol 27:147–154

167. Wang Z, Leung KMY, Sung Y-H, Dudgeon D, Qiu J-W (2021) Recovery of tropical marine benthos after a trawl ban demonstrates linkage between abiotic and biotic changes. Commun Biol 4:212

168. Wen C, Pratchett MS, Shao K, Kan K, Chan BKK (2010) Effects of habitat modification on coastal fish assemblages. J Fish Biol 77:1674–1687

169. Wenger AS, McCormick MI, McLeod IM, Jones GP (2013) Suspended sediment alters predator– prey interactions between two coral reef fishes. Coral Reefs 32:369–374

170. West KM, Stat M, Harvey ES, Skepper CL, DiBattista JD, Richards ZT, Travers MJ, Newman SJ, Bunce M (2020) eDNA metabarcoding survey reveals fine-scale coral reef community variation across a remote, tropical island ecosystem. Mol Ecol 29:1069–1086

171. Whittaker RH (1972) Evolution and measurement of species diversity. Taxon 21:213–251

172. Williams GA, Chan BKK, Dong Y (2019) Rocky Shores of Mainland China, Taiwan and Hong Kong. Interactions in the Marine Benthos

173. Williamson DH, Ceccarelli DM, Evans RD, Jones GP, Russ GR (2014) Habitat dynamics, marine reserve status, and the decline and recovery of coral reef fish communities. Ecol Evol 4:337– 354

174. Wilson SK, Fulton CJ, Depczynski M, Holmes TH, Noble MM, Radford B, Tinkler P (2014) Seasonal changes in habitat structure underpin shifts in macroalgae-associated tropical fish communities. Mar Biol 161:2597–2607

175. Yamamoto S, Masuda R, Sato Y, Sado T, Araki H, Kondoh M, Minamoto T, Miya M (2017) Environmental DNA metabarcoding reveals local fish communities in a species-rich coastal sea. Sci Rep 7:40368

176. Yates F (1934) The analysis of multiple classifications with unequal numbers in the different classes. J Am Stat Assoc 29:51–66

177. Yeung YH, Xie JY, Kwok CK, Kei K, Ang P, Chan LL, Dellisanti W, Cheang CC, Chow WK, Qiu JW (2021) Hong Kong’s subtropical scleractinian coral communities: Baseline, environmental drivers and management implications. Mar Pollut Bull 167:112289

178. Yiu SKF, Chow CFY, Tsang SHT, Zhang X, Chung JTH, Sin SYT, Chow WK, Chan LL (2022) New record of the Japanese Seahorse Hippocampus mohnikei Bleeker, 1853 (Syngnathiformes: Syngnathidae) in Hong Kong waters. Check List 18:455–461

179. Yung Y-K, Wong CK, Yau K, Qian PY (2001) Long-term changes in water quality and phytoplankton characteristics in Port Shelter, Hong Kong, from 1988–1998. Mar Pollut Bull 42:981–992

180. Zamani NP, Zuhdi MF, Madduppa H (2022) Environmental DNA biomonitoring reveals seasonal patterns in coral reef fish community structure. Environ Biol Fishes 105:971–991

181. Zhu M, Kuroki M, Kobayashi T, Yamakawa T, Sado T, Kodama K, Horiguchi T, Miya M (2023) Comparison of fish fauna evaluated using aqueous eDNA, sedimentary eDNA, and catch surveys in Tokyo Bay, Central Japan. Journal of Marine Systems 240:

182. Zou K, Chen J, Ruan H, Li Z, Guo W, Li M, Liu L (2020) eDNA metabarcoding as a promising conservation tool for monitoring fish diversity in a coastal wetland of the Pearl River Estuary compared to bottom trawling. Science of the Total Environment 702:

