## Supplementary Material for "Characterisation of coastal reef fish assemblages across an estuarine-urbanisation gradient using underwater visual survey and environmental DNA metabarcoding"

### Supplementary Materials

***Supplementary Figures***

**Supplementary Figure 1.** Non-metric multidimensional scaling (nMDS) plot of fish assemblages based on Jaccard's distances of presence-absence data from underwater visual census and eDNA metabarcoding from both sampling months. Ellipses are drawn based on 95% confidence. The size of data points reflects fish generic richness. UVC: underwater visual census (olive), eDNA: eDNA metabarcoding (navy), Lantau (circle), S-HK: South Hong Kong (square), SE-HK: Southeast Hong Kong (diamond), E-SK: East Sai Kung (triangle), N-SK: North Sai Kung (upside-down triangle).

**Supplementary Figure 2.** Five-dimensional Venn diagrams showing the number of fish genera that were able to retrieve using (a) underwater visual census or (b) eDNA metabarcoding across five regions. Lantau (yellow), S-HK: South Hong Kong (green), SE-HK: Southeast Hong Kong (black), E-SK: East Sai Kung (sky blue), N-SK: North Sai Kung (pink).

**Supplementary Figure 3.** Two-dimensional Venn diagrams showing the number of fish genera that were able to retrieve using (a) underwater visual census or (b) eDNA metabarcoding between two sampling months. Dry: November/December (gray), Wet: May/June (dark red).

**Supplementary Figure 4.** Images of potential novel fish species to the waters of Hong Kong, including a) *Enneapterygius cf. hsiojenae*, b) *Parascorpaena cf. aurita*, c) *Cryptocentrus cf. cinctus*, d) *Cryptocentrus cf. leucostictus*, and e) *Cryptocentrus cf. melanopus*. Photo images (a) and (b) were taken by YDP from Long Ke Wan in May 2020 for *E. hsiojenae* and in June 2019 for *P. aurita*, and later verified by fish biologists Dr. Chen-Lu Lee and Dr. Hiroyuki Motomura through personal communication with YDP. The rest of photo images (c, d, e) were taken by JH during the visual surveys of this study.

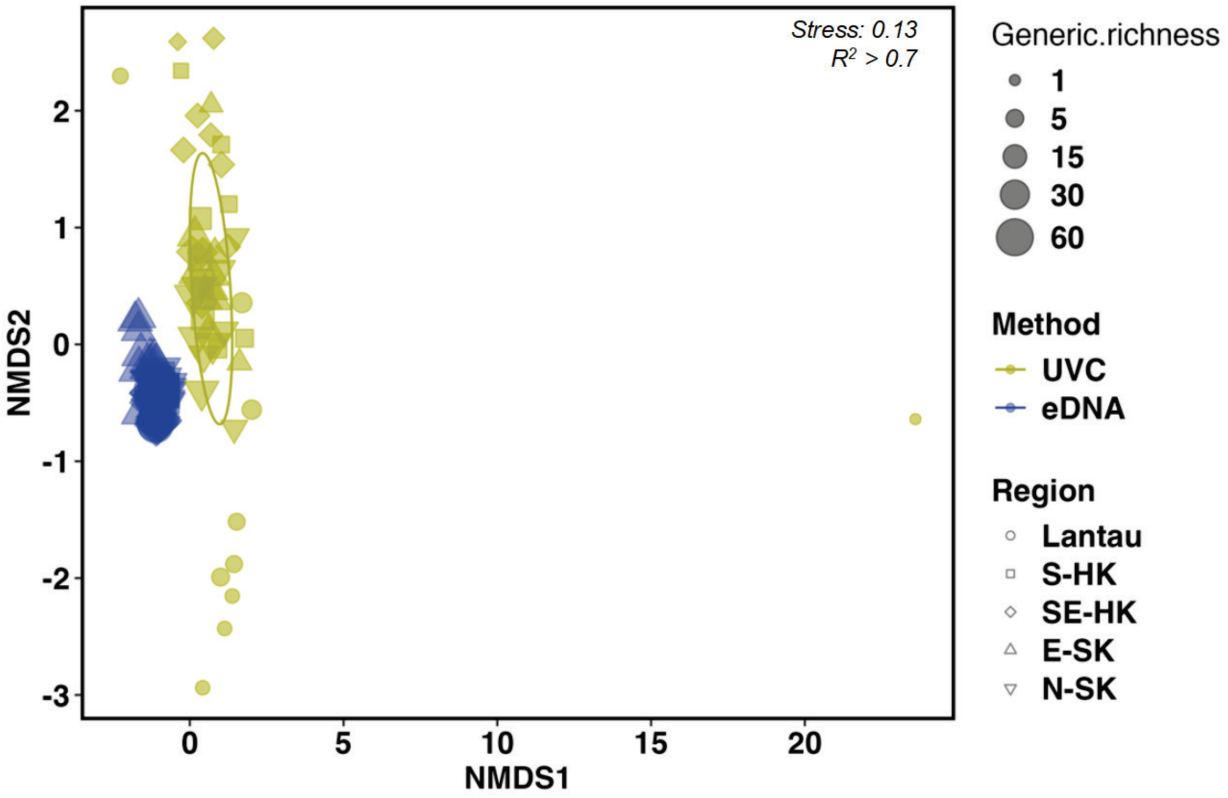


Supplementary Figure 1. Non-metric multidimensional scaling (nMDS) plot of fish assemblages based on Jaccard's distances of presence-absence data from underwater visual census and eDNA metabarcoding from both sampling months. Ellipses are drawn based on 95% confidence. The size of data points reflects fish generic richness. UVC: underwater visual census (olive), eDNA: eDNA metabarcoding (navy), Lantau (circle), S-HK: South Hong Kong (square), SE-HK: Southeast Hong Kong (diamond), E-SK: East Sai Kung (triangle), N-SK: North Sai Kung (upside-down triangle).

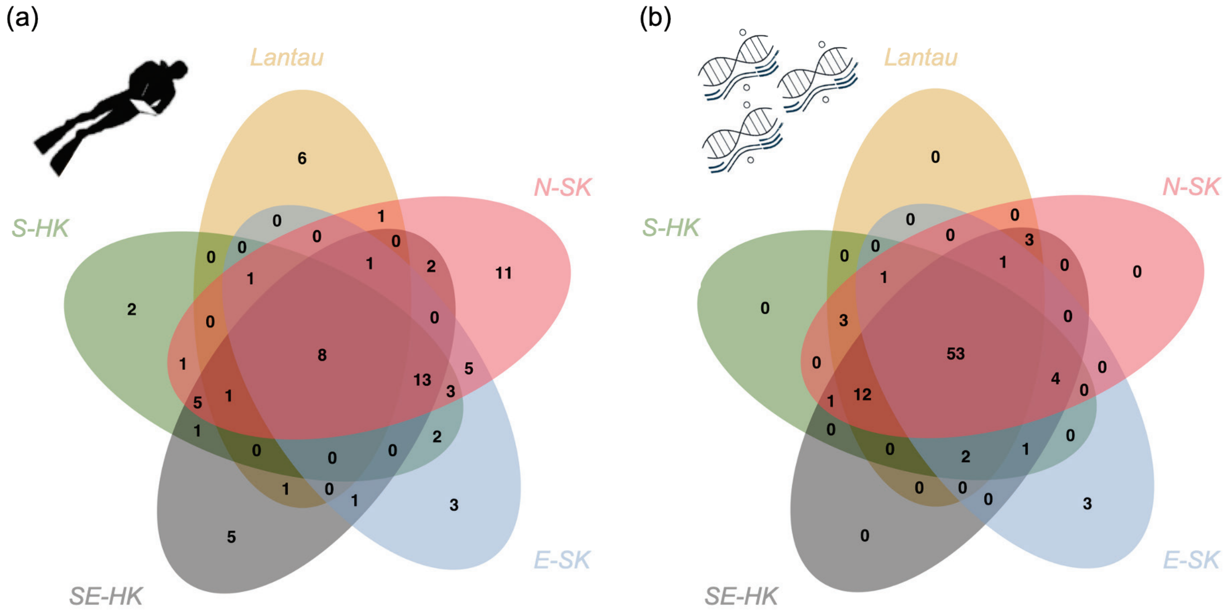


Supplementary Figure 2. Five-dimensional Venn diagrams showing the number of fish genera that were able to retrieve using (a) underwater visual census or (b) eDNA metabarcoding across five regions. Lantau (yellow), S-HK: South Hong Kong (green), SE-HK: Southeast Hong Kong (black), E-SK: East Sai Kung (sky blue), N-SK: North Sai Kung (pink).

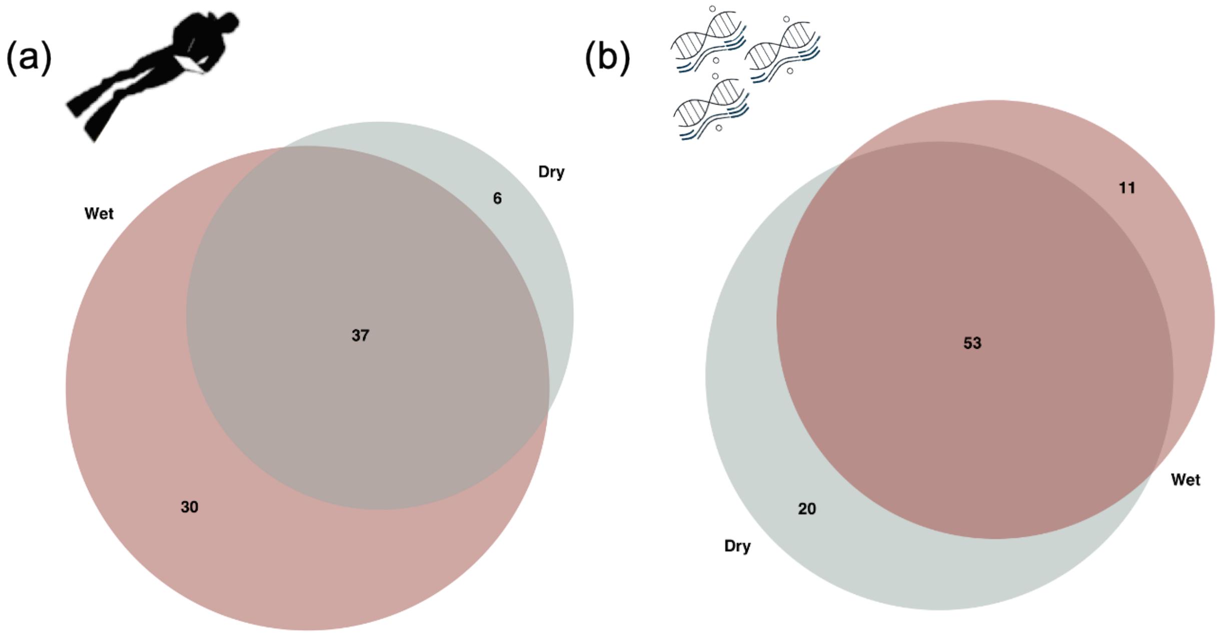


Supplementary Figure 3. Two-dimensional Venn diagrams showing the number of fish genera that were able to retrieve using (a) underwater visual census or (b) eDNA metabarcoding between two sampling months. Wet: May/June (dark red), Dry: November/December (gray).

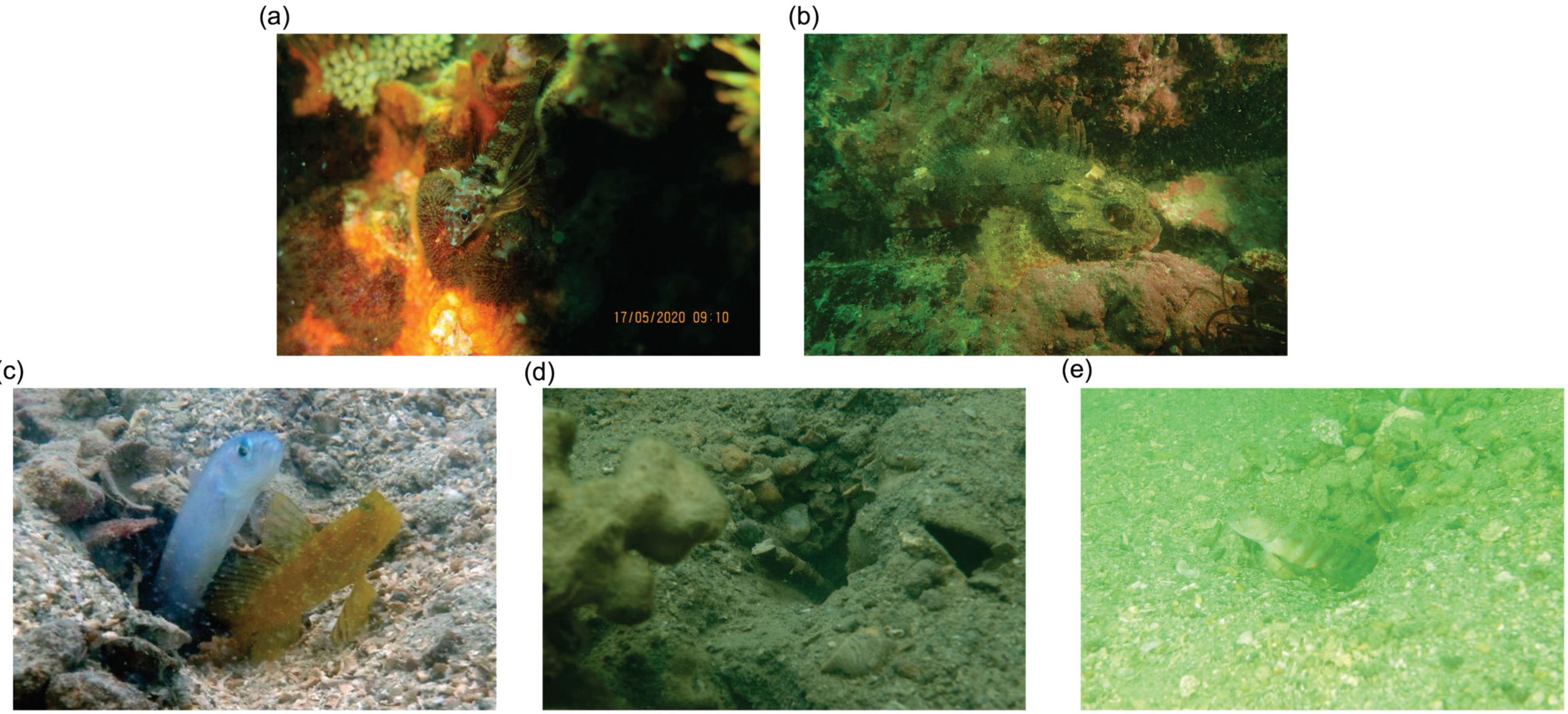


Supplementary Figure 4. Images of potential novel fish species to the waters of Hong Kong, including a) *Enneapterygius cf. hsiojenae*, b) *Parascorpaena cf. aurita*, c) *Cryptocentrus cf. cinctus* (right), d) *Cryptocentrus cf. leucostictus*, and e) *Cryptocentrus cf. melanopus*. Photo images (a) and (b) were taken by YDP from Long Ke Wan in May 2020 for *E. hsiojenae* and in June 2019 for *P. aurita*, and later verified by fish biologists Dr. Chen-Lu Lee and Dr. Hiroyuki Motomura through personal communication with YDP. The rest of photo images (c, d, e) were taken by JH during the visual surveys of this study.

*Supplementary tables*

**Supplementary Table 1.** The GPS information for each site and their respective region. S-HK: South Hong Kong, SE-HK: Southeast Hong Kong, E-SK: East Sai Kung, N-SK: North Sai Kung.

**Supplementary Table 2.** An amalgamation of fish species recorded during underwater UVC and detected by eDNA metabarcoding from water samples taken from surface level. Species that were recorded or detected were given a tick (v) while the otherwise were left blank.

**Supplementary Table 3.** Information regarding the general water column position, habitat association, diet, and preferred aquatic environment, as well as the presence and absence record of recorded fish genera using underwater visual census (UVC) or eDNA metabarcoding (eDNA) across the five regions of Hong Kong. Genera that were recorded or detected were given a tick (v) while the otherwise were left blank. S-HK: South Hong Kong, SE-HK: Southeast Hong Kong, E-SK: East Sai Kung, N-SK: North Sai Kung.

**Supplementary Table 4.** Global and regional differences in the (a) total number and (b) average and standard deviation (Std.) of fish genera recorded between regions across both and separate sampling seasons using UVC and eDNA metabarcoding.

**Supplementary Table 5.** Permutational Multivariate Analysis of Variance (PERMANOVA) and PERMDISP tests for pairwise comparisons of fish composition among regions for both sampling season combined and during either season for (a) UVC and (b) eDNA metabarcoding. df: degree of freedom; Significant results are in bold. ***: $p < 0.001$; **: $p < 0.01$; *: $p < 0.05$.

**Supplementary Table 6.** Pairwise comparisons for Permutational Multivariate Analysis of Variance (PERMANOVA) – *PairwiseAdonis* and for PERMDISP – *Betadisper* among regions for both sampling seasons combined and for only wet and dry seasons for (a-c) UVC and (d-f) eDNA metabarcoding. S-HK: South Hong Kong, SE-HK: Southeast Hong Kong, E-SK: East Sai Kung, N-SK: North Sai Kung. Significant results are presented as ***: $p < 0.001$; **: $p < 0.01$; *: $p < 0.05$.

**Supplementary Table 7.** Indicator species analysis (ISA) results for the combinations of region and sampling seasons based on underwater visual census data. Significant results are presented as ***: $p < 0.001$; **: $p < 0.01$; *: $p < 0.05$.

**Supplementary Table 8.** Indicator species analysis (ISA) results for the combinations of region and sampling seasons based on eDNA metabarcoding data. Significant results are presented as ***: $p < 0.001$; **: $p < 0.01$; *: $p < 0.05$.

Supplementary Table 1. The GPS information for each site and their respective region. S-HK: South Hong Kong, SE-HK: Southeast Hong Kong, E-SK: East Sai Kung, N-SK: North Sai Kung.

| Site | Region | Northing | Easting |
|---|---|---|---|
| Sam Pak Wan | Lantau | 22°18'32.22"N | 114° 1'22.07"E |
| Peng Chau | Lantau | 22°17'14.17"N | 114° 2'38.15"E |
| Hei Ling Chau | Lantau | 22°15'30.55"N | 114° 2'20.44"E |
| Chi Ma Wan | Lantau | 22°14'20.62"N | 114° 0'4.29"E |
| Cheung Chau | Lantau | 22°12'57.04"N | 114° 1'29.48"E |
| Po Toi | S-HK | 22° 9'34.99"N | 114°15'26.67"E |
| Beaufort Island | S-HK | 22°10'52.14"N | 114°14'47.25"E |
| Cape D'Aguilar | S-HK | 22°12'24.94"N | 114°15'21.46"E |
| Stanley Village | S-HK | 22°12'56.20"N | 114°12'45.83"E |
| Sham Wan | S-HK | 22°11'16.79"N | 114° 8'5.43"E |
| Ninepin South | SE-HK | 22°15'30.70"N | 114°21'2.72"E |
| Ninepin North | SE-HK | 22°16'18.78"N | 114°20'55.69"E |
| Trio Rocks | SE-HK | 22°18'4.26"N | 114°19'7.85"E |
| Steep Island | SE-HK | 22°16'25.59"N | 114°18'45.22"E |
| Tung Lung Chau | SE-HK | 22°14'26.51"N | 114°17'8.24"E |
| Long Ke Wan | E-SK | 22°22'28.57"N | 114°22'55.06"E |
| Bluff Island | E-SK | 22°19'30.52"N | 114°21'13.51"E |
| Tai She Wan | E-SK | 22°21'25.93"N | 114°20'22.65"E |
| Shelter Island | E-SK | 22°19'49.69"N | 114°17'33.72"E |
| Sharp Island | E-SK | 22°21'40.51"N | 114°17'30.31"E |
| Tung Ping Chau | N-SK | 22°32'48.10"N | 114°26'11.3"E |
| Kat O | N-SK | 22°33'4.40"N | 114°18'35.88"E |
| Crescent Island | N-SK | 22°32'15.92"N | 114°19'8.38"E |
| Sai Lau Kong | N-SK | 22°31'24.32"N | 114°17'9.03"E |
| Gruff Head | N-SK | 22°28'30.61"N | 114°19'25.24"E |

Supplementary Table 2. An amalgamation of fish species recorded using underwater visual census (UVC) or eDNA metabarcoding (eDNA) from water samples taken from surface level. Species that were recorded or detected were given a tick (v) while the otherwise were left blank. Species in bold and red represent species that have not been officially reported from Hong Kong.

| Family | Genus | Species | UVC | eDNA |
|---|---|---|---|---|
| *Pomacentridae* | *Abudefduf* | *Abudefduf.bengalensis* | ✓ | ✓ |
| *Pomacentridae* | *Abudefduf* | *Abudefduf.sordidus* | ✓ | ✓ |
| *Pomacentridae* | *Abudefduf* | *Abudefduf.vaigiensis* | ✓ | ✓ |
| *Sparidae* | *Acanthopagrus* | *Acanthopagrus.latus* | ✓ | |
| *Sparidae* | *Acanthopagrus* | *Acanthopagrus.pacificus* | | ✓ |
| *Carangidae* | *Alectis* | *Alectis.indica* | | ✓ |
| *Carangidae* | *Alepes* | *Alepes_spp.1* | | ✓ |
| *Carangidae* | *Alepes* | *Alepes.djedaba* | | ✓ |
| *Ambassidae* | *Ambassis* | *Ambassis_spp.1* | | ✓ |
| *Ambassidae* | *Ambassis* | *Ambassis.sp.* | ✓ | |
| *Gobiidae* | *Amblyeleotris* | *Amblyeleotris.gymnocephala* | ✓ | |
| *Gobiidae* | *Amblyeleotris* | *Amblyeleotris.japonica* | ✓ | |
| *Gobiidae* | *Amblyeleotris* | *Amblyeleotris.periophthalma* | ✓ | |
| *Gobiidae* | *Amblygobius* | *Amblygobius.phalaena* | ✓ | ✓ |
| *Pomacentridae* | *Amphiprion* | *Amphiprion.clarkii* | ✓ | ✓ |
| *Apogonidae* | *Apogonichthyoides* | *Apogonichthyoides.cathetogramma* | ✓ | |
| *Tetraodontidae* | *Arothron* | *Arothron.hispidus* | ✓ | ✓ |
| *Gobiidae* | *Asterropteryx* | *Asterropteryx.semipunctata* | ✓ | ✓ |
| *Atherinidae* | *Atherinomorus* | *Atherinomorus.pinguis* | | ✓ |
| *Atherinidae* | *Atherinomorus* | *Atherinomorus.sp.* | ✓ | |
| ***Gobiidae*** | ***Bathygobius*** | ***Bathygobius.cotticeps*** | | ✓ |
| *Gobiidae* | *Bathygobius* | *Bathygobius.hongkongensis* | | ✓ |
| *Gobiidae* | *Bathygobius* | *Bathygobius.sp.* | ✓ | |
| *Blenniidae* | | *Blenniidae.sp.* | ✓ | |
| *Bothidae* | | *Bothidae.sp.* | ✓ | |
| *Callionymidae* | | *Callionymidae.sp.* | ✓ | |
| *Callionymidae* | *Callionymus* | *Callionymus.curvicornis* | | ✓ |
| ***Carangidae*** | ***Carangoides*** | ***Carangoides.hedlandensis*** | | ✓ |
| *Carangidae* | *Carangoides* | *Carangoides.praeustus* | ✓ | |
| *Epinephelidae* | *Cephalopholis* | *Cephalopholis.boenak* | ✓ | ✓ |
| *Chaetodontidae* | *Chaetodon* | *Chaetodon.auripes* | | ✓ |
| *Chaetodontidae* | *Chaetodon* | *Chaetodon.speculum* | ✓ | |
| *Chaetodontidae* | *Chaetodon* | *Chaetodon.wiebeli* | ✓ | |
| *Labridae* | *Cheilinus* | *Cheilinus.chlorourus* | ✓ | |
| *Apogonidae* | *Cheilodipterus* | *Cheilodipterus.intermedius* | ✓ | |
| *Pomacentridae* | *Chromis* | *Chromis_spp.1* | | ✓ |
| *Pomacentridae* | *Chromis* | *Chromis.fumea* | ✓ | |
| *Pomacentridae* | *Chromis* | *Chromis.notata* | ✓ | |
| *Cichlidae* | | *Cichlidae_sp.1* | | ✓ |
| *Dorosomatidae* | *Clupanodon* | *Clupanodon.thrissa* | | ✓ |
| *Mugilidae* | *Crenimugil* | *Crenimugil.crenilabis* | ✓ | |
| ***Mugilidae*** | ***Crenimugil*** | ***Crenimugil.pedaraki*** | | ✓ |
| *Mugilidae* | *Crenimugil* | *Crenimugil.seheli* | | ✓ |
| ***Gobiidae*** | ***Cryptocentrus*** | ***Cryptocentrus.albidorsus*** | ✓ | |
| ***Gobiidae*** | ***Cryptocentrus*** | ***Cryptocentrus.caeruleomaculatus*** | ✓ | ✓ |
| *Gobiidae* | *Cryptocentrus* | *Cryptocentrus.cinctus* | ✓ | |
| ***Gobiidae*** | ***Cryptocentrus*** | ***Cryptocentrus.leucostictus*** | ✓ | |
| ***Gobiidae*** | ***Cryptocentrus*** | ***Cryptocentrus.melanopus*** | ✓ | |

| | | | | |
|---|---|---|---|---|
| **Gobiidae** | **Cryptocentrus** | **Cryptocentrus.sericus** | **✓** | |
| Gobiidae | Cryptocentrus | Cryptocentrus.strigilliceps | ✓ | |
| Pomacentridae | Dascyllus | Dascyllus.trimaculatus | ✓ | |
| Carangidae | Decapterus | Decapterus.maruadsi | | ✓ |
| Scorpaenidae | Dendrochirus | Dendrochirus.zebra | ✓ | |
| **Leiognathidae** | **Deveximentum** | **Deveximentum.indicium** | | **✓** |
| Haemulidae | Diagramma | Diagramma.pictum | ✓ | |
| Haemulidae | Diagramma | Diagramma_spp.1 | | ✓ |
| Liopropomatidae | Diploprion | Diploprion.bifasciatum | ✓ | |
| Drepaneidae | Drepane | Drepane_spp.1 | | ✓ |
| Polynemidae | Eleutheronema | Eleutheronema.tetradactylum | | ✓ |
| Elopidae | Elops | Elops_spp.1 | | ✓ |
| Engraulidae | Encrasicholina | Encrasicholina_spp.1 | | ✓ |
| Engraulidae | Encrasicholina | Encrasicholina.heteroloba | | ✓ |
| Engraulidae | Encrasicholina | Encrasicholina.punctifer | | ✓ |
| Tripterygiidae | Enneapterygius | Enneapterygius.etheostoma | ✓ | ✓ |
| **Tripterygiidae** | **Enneapterygius** | **Enneapterygius.hsiojenae** | | **✓** |
| Blenniidae | Entomacrodus | Entomacrodus.stellifer | | ✓ |
| Epinephelidae | Epinephelus | Epinephelus_spp.1 | | ✓ |
| Epinephelidae | Epinephelus | Epinephelus.bleekeri | ✓ | |
| Epinephelidae | Epinephelus | Epinephelus.fasciatomaculosus | ✓ | |
| Epinephelidae | Epinephelus | Epinephelus.quoyanus | ✓ | |
| **Leiognathidae** | **Equulites** | **Equulites.oblongus** | | **✓** |
| Scombridae | Euthynnus | Euthynnus_spp.1 | | ✓ |
| Gobiidae | Eviota | Eviota.storthynx | ✓ | |
| Fistulariidae | Fistularia | Fistularia.commersonii | ✓ | |
| Leiognathidae | Gazza | Gazza.minuta | | ✓ |
| Gerreidae | Gerres | Gerres_spp.1 | | ✓ |
| Gerreidae | Gerres | Gerres_spp.2 | | ✓ |
| Gerreidae | Gerres | Gerres_spp.3 | | ✓ |
| Gerreidae | Gerres | Gerres_spp.4 | | ✓ |
| Gerreidae | Gerres | Gerres.filamentosus | ✓ | |
| Gerreidae | Gerres | Gerres.oblongus | ✓ | |
| Gerreidae | Gerres | Gerres.oyena | ✓ | |
| Gobiidae | | Gobiidae.sp. | ✓ | |
| Gobiidae | | Gobiidae.sp..1 | ✓ | |
| Gobiidae | | Gobiidae.sp..2 | ✓ | |
| Gobiidae | | Gobiidae.sp..3 | ✓ | |
| Gobiidae | | Gobiidae.sp..4 | ✓ | |
| Gobiidae | | Gobiidae.sp..5 | ✓ | |
| Muraenidae | Gymnothorax | Gymnothorax_spp.1 | | ✓ |
| Muraenidae | Gymnothorax | Gymnothorax.sp. | ✓ | |
| Labridae | Halichoeres | Halichoeres.hartzfeldii | ✓ | |
| Labridae | Halichoeres | Halichoeres.kneri | ✓ | |
| Labridae | Halichoeres | Halichoeres.marginatus | ✓ | |
| Labridae | Halichoeres | Halichoeres.nebulosus | ✓ | |
| Labridae | Halichoeres | Halichoeres.nigrescens | ✓ | |
| Dorosomatidae | Hilsa | Hilsa.kelee | | ✓ |
| **Atherinidae** | **Hypoatherina** | **Hypoatherina.woodwardi** | | **✓** |
| Hemiramphidae | Hyporhamphus | Hyporhamphus.quoyi | | ✓ |
| Gobiidae | Istigobius | Istigobius.campbelli | ✓ | ✓ |
| Gobiidae | Istigobius | Istigobius.decoratus | ✓ | |
| Gobiidae | Istigobius | Istigobius.sp. | ✓ | |
| Dorosomatidae | Konosirus | Konosirus.punctatus | | ✓ |

| | | | | |
|---|---|---|---|---|
| Labridae | | Labridae_sp.1 | | √ |
| Labridae | | Labridae.sp. | √ | |
| Tetraodontidae | Lagocephalus | Lagocephalus.spadiceus | | √ |
| Sciaenidae | Larimichthys | Larimichthys.crocea | | √ |
| Leiognathidae | | Leiognathidae_sp.1 | | √ |
| Leiognathidae | | Leiognathidae_sp.2 | | √ |
| Trichiuridae | Lepturacanthus | Lepturacanthus.savala | | √ |
| Lutjanidae | Lutjanus | Lutjanus_spp.1 | | √ |
| Lutjanidae | Lutjanus | Lutjanus.russellii | √ | √ |
| Microcanthidae | Microcanthus | Microcanthus.strigatus | √ | |
| Monacanthidae | Monacanthus | Monacanthus.chinensis | √ | √ |
| Mugilidae | Mugil | Mugil.cephalus | | √ |
| Mugilidae | | Mugilidae_sp.1 | | √ |
| Mugilidae | | Mugilidae_sp.2 | | √ |
| Gobiidae | Myersina | Myersina.filifer | √ | |
| Dorosomatidae | Nematalosa | Nematalosa.japonica | | √ |
| Dorosomatidae | Nematalosa | Nematalosa.nasus | | √ |
| **Pomacentridae** | **Neopomacentrus** | **Neopomacentrus.anabatoides** | | **√** |
| Pomacentridae | Neopomacentrus | Neopomacentrus.bankieri | √ | |
| Pomacentridae | Neopomacentrus | Neopomacentrus.cyanomos | √ | √ |
| Sciaenidae | Nibea | Nibea.albiflora | | √ |
| Leiognathidae | Nuchequula | Nuchequula.nuchalis | | √ |
| Blenniidae | Omobranchus | Omobranchus.sp. | √ | |
| Apogonidae | Ostorhinchus | Ostorhinchus.cookii | | √ |
| Apogonidae | Ostorhinchus | Ostorhinchus.doederleini | √ | √ |
| Apogonidae | Ostorhinchus | Ostorhinchus.fasciatus | √ | |
| Apogonidae | Ostorhinchus | Ostorhinchus.fleurieu | √ | |
| Apogonidae | Ostorhinchus | Ostorhinchus.holotaenia | √ | |
| Ostraciidae | Ostracion | Ostracion.cubicum | √ | |
| Gobiidae | Oxyurichthys | Oxyurichthys.auchenolepis | | √ |
| Sparidae | Pagrus | Pagrus.major | √ | |
| Tetrarogidae | Paracentropogon | Paracentropogon.longispinis | √ | |
| Monacanthidae | Paramonacanthus | Paramonacanthus.sulcatus | √ | |
| Pinguipedidae | Parapercis | Parapercis.snyderi | √ | |
| Haemulidae | Parapristipoma | Parapristipoma.trilineatum | | √ |
| **Scorpaenidae** | **Parascorpaena** | **Parascorpaena.aurita** | | **√** |
| Scorpaenidae | Parascorpaena | Parascorpaena.mossambica | | √ |
| Scorpaenidae | Parascorpaena | Parascorpaena.sp. | √ | |
| **Exocoetidae** | **Parexocoetus** | **Parexocoetus.mento** | | **√** |
| Microdesmidae | Parioglossus | Parioglossus.dotui | √ | |
| Mullidae | Parupeneus | Parupeneus.biaculeatus | √ | √ |
| Mullidae | Parupeneus | Parupeneus.ciliatus | √ | |
| Mullidae | Parupeneus | Parupeneus.indicus | √ | √ |
| Terapontidae | Pelates | Pelates.quadrilineatus | √ | |
| Pempheridae | Pempheris | Pempheris.oualensis | √ | |
| Pempheridae | Pempheris | Pempheris.schwenkii | √ | √ |
| Sciaenidae | Pennahia | Pennahia.aneus | | √ |
| Blenniidae | Petroscirtes | Petroscirtes.breviceps | √ | √ |
| Leiognathidae | Photopectoralis | Photopectoralis.bindus | | √ |
| Mugilidae | Planiliza | Planiliza.macrolepis | | √ |
| Mugilidae | Planiliza | Planiliza.sp. | √ | |
| Mugilidae | Planiliza | Planiliza.subviridis | | √ |
| Haemulidae | Plectorhinchus | Plectorhinchus.gibbosus | | √ |
| | | Pleuronectiformes_f_sp.1 | | √ |

| | | | | |
|---|---|---|---|---|
| *Mugilidae* | *Plicomugil* | *Plicomugil.labiosus* | | √ |
| *Plotosidae* | *Plotosus* | *Plotosus_spp.1* | | √ |
| *Plotosidae* | *Plotosus* | *Plotosus.lineatus* | √ | |
| *Pomacentridae* | *Pomacentrus* | *Pomacentrus.chrysurus* | √ | |
| *Labridae* | *Pteragogus* | *Pteragogus.enneacanthus* | √ | |
| *Microdesmidae* | *Ptereleotris* | *Ptereleotris.hanae* | √ | |
| *Microdesmidae* | *Ptereleotris* | *Ptereleotris.microlepis* | √ | |
| *Scombridae* | *Rastrelliger* | *Rastrelliger.kanagurta* | | √ |
| *Sparidae* | *Rhabdosargus* | *Rhabdosargus.sarba* | √ | √ |
| *Dorosomatidae* | *Sardinella* | *Sardinella_spp.1* | | √ |
| *Dorosomatidae* | *Sardinella* | *Sardinella_spp.2* | | √ |
| *Dorosomatidae* | *Sardinella* | *Sardinella_spp.3* | | √ |
| *Dorosomatidae* | *Sardinella* | *Sardinella_spp.4* | | √ |
| *Scaridae* | *Scarus* | *Scarus.ghobban* | √ | |
| *Scaridae* | *Scarus* | *Scarus.sp..1* | √ | |
| *Scaridae* | *Scarus* | *Scarus.sp..2* | √ | |
| *Scatophagidae* | *Scatophagus* | *Scatophagus.argus* | √ | |
| *Sciaenidae* | *Sciaenops* | *Sciaenops.ocellatus* | | √ |
| *Scorpaenidae* | *Scorpaenopsis* | *Scorpaenopsis.cirrosa* | √ | |
| *Sebastidae* | *Sebastiscus* | *Sebastiscus.marmoratus* | √ | √ |
| *Carangidae* | *Selar* | *Selar.crumenophthalmus* | | √ |
| *Carangidae* | *Selaroides* | *Selaroides.leptolepis* | √ | √ |
| *Siganidae* | *Siganus* | *Siganus.fuscescens* | √ | √ |
| *Sillaginidae* | *Sillago* | *Sillago.aeolus* | √ | √ |
| *Sillaginidae* | *Sillago* | *Sillago.asiatica* | | √ |
| *Sparidae* | | *Sparidae_sp.1* | | √ |
| *Sparidae* | | *Sparidae_sp.2* | | √ |
| *Sphyraenidae* | *Sphyraena* | *Sphyraena_spp.1* | | √ |
| *Sphyraenidae* | *Sphyraena* | *Sphyraena.jello* | | √ |
| *Spratelloididae* | *Spratelloides* | *Spratelloides.gracilis* | | √ |
| *Labridae* | *Stethojulis* | *Stethojulis.terina* | √ | √ |
| *Engraulidae* | *Stolephorus* | *Stolephorus_spp.1* | | √ |
| *Synodontidae* | *Synodus* | *Synodus.dermatogenys* | √ | |
| *Synodontidae* | *Synodus* | *Synodus.sp.* | √ | |
| *Apogonidae* | *Taeniamia* | *Taeniamia_spp.1* | | √ |
| *Apogonidae* | *Taeniamia* | *Taeniamia.fucata* | √ | |
| *Tetraodontidae* | *Takifugu* | *Takifugu_spp.1* | | √ |
| *Tetraodontidae* | *Takifugu* | *Takifugu.flavipterus* | √ | |
| *Labridae* | *Thalassoma* | *Thalassoma_spp.1* | | √ |
| *Labridae* | *Thalassoma* | *Thalassoma.lunare* | √ | |
| *Engraulidae* | *Thryssa* | *Thryssa_spp.1* | | √ |
| *Engraulidae* | *Thryssa* | *Thryssa.kammalensis* | | √ |
| *Platycephalidae* | *Thysanophrys* | *Thysanophrys.celebica* | √ | |
| *Carangidae* | *Trachinotus* | *Trachinotus_spp.1* | | √ |
| *Trichonotidae* | *Trichonotus* | *Trichonotus.setiger* | √ | |
| *Gobiidae* | *Tridentiger* | *Tridentiger.trigonocephalus* | √ | √ |
| *Belonidae* | *Tylosurus* | *Tylosurus_spp.1* | | √ |
| *Mullidae* | *Upeneus* | *Upeneus.sulphureus* | | √ |
| *Mullidae* | *Upeneus* | *Upeneus.tragula* | √ | √ |
| *Gobiidae* | *Valenciennea* | *Valenciennea.immaculata* | √ | |
| *Gobiidae* | *Valenciennea* | *Valenciennea.muralis* | √ | |
| *Zanclidae* | *Zanclus* | *Zanclus.cornutus* | √ | |

Supplementary Table 3. The information regarding the water column position, habitat association, diet, and preferred aquatic environment (environment), as well as the presence and absence record of recorded fish genera using underwater visual census (UVC) or eDNA metabarcoding (eDNA) across the five regions of Hong Kong. Genera that were recorded or detected were given a tick (v) while the otherwise were left blank. S-HK: South Hong Kong, SE-HK: Southeast Hong Kong, E-SK: East Sai Kung, N-SK: North Sai Kung.

| | | | | | UVC | | | | | eDNA | | | | |
|---|---|---|---|---|---|---|---|---|---|---|---|---|---|---|
| Genus | Water column position | Habitat association | Diet | Environment | Lantau | S-HK | SE-HK | E-SK | N-SK | Lantau | S-HK | SE-HK | E-SK | N-SK |
| *Abudefduf* | Benthopelagic | Reef-associated | Omnivorous | Marine/Brackish | | ✓ | ✓ | ✓ | ✓ | ✓ | ✓ | ✓ | ✓ | ✓ |
| *Acanthopagrus* | Benthopelagic | Demersal | Invertivorous | Marine/Brackish/Freshwater | ✓ | | ✓ | | | ✓ | ✓ | ✓ | | ✓ |
| *Alectis* | Benthopelagic | Reef-associated | Invertivorous.Piscivorous | Marine/Brackish | | | | | | ✓ | ✓ | ✓ | | ✓ |
| *Alepes* | Benthopelagic | Reef-associated | Invertivorous.Piscivorous | Marine/Brackish | | | | | | ✓ | ✓ | ✓ | ✓ | ✓ |
| *Ambassis* | Benthopelagic | Reef-associated | Invertivorous | Marine | | | | | ✓ | ✓ | ✓ | ✓ | ✓ | ✓ |
| *Amblyeleotris* | Benthic | Reef-associated | Omnivorous | Marine/Brackish | | ✓ | ✓ | | ✓ | | | | | |
| *Amblygobius* | Benthic | Reef-associated | Omnivorous | Marine | | | | ✓ | ✓ | ✓ | ✓ | | ✓ | ✓ |
| *Amphiprion* | Benthopelagic | Reef-associated | Omnivorous | Marine | | ✓ | ✓ | ✓ | ✓ | ✓ | ✓ | ✓ | ✓ | ✓ |
| *Apogonichthyoides* | Benthopelagic | Reef-associated | Invertivorous | Marine | | ✓ | ✓ | ✓ | ✓ | | | | | |
| *Arothron* | Benthopelagic | Reef-associated | Omnivorous | Marine/Brackish | | | ✓ | | | ✓ | ✓ | ✓ | | ✓ |
| *Asterropteryx* | Benthic | Reef-associated | Invertivorous | Marine | | | | ✓ | ✓ | ✓ | ✓ | | | ✓ |
| *Atherinomorus* | Benthopelagic | Reef-associated | Planktivorous | Marine/Brackish/Freshwater | | | | | ✓ | ✓ | ✓ | ✓ | ✓ | ✓ |
| *Bathygobius* | Benthic | Reef-associated | Invertivorous | Marine/Brackish | ✓ | | | | ✓ | ✓ | ✓ | ✓ | ✓ | ✓ |
| *Callionymus* | Benthic | Demersal | Invertivorous | Marine/Brackish/Freshwater | | | | | | ✓ | ✓ | ✓ | | ✓ |
| *Carangoides* | Benthopelagic | Reef-associated/Demersal | Invertivorous | Marine | | | | | ✓ | ✓ | ✓ | ✓ | ✓ | ✓ |
| *Cephalopholis* | Benthopelagic | Reef-associated | Invertivorous.Piscivorous | Marine | | ✓ | ✓ | ✓ | ✓ | ✓ | ✓ | ✓ | ✓ | ✓ |
| *Chaetodon* | Benthopelagic | Reef-associated | Invertivorous/Omnivorous | Marine | | | ✓ | | ✓ | ✓ | ✓ | ✓ | ✓ | |
| *Cheilinus* | Benthopelagic | Reef-associated | Invertivorous | Marine | | | | ✓ | ✓ | | | | | |
| *Cheilodipterus* | Benthopelagic | Reef-associated | Invertivorous | Marine | | ✓ | | ✓ | ✓ | | | | | |
| *Chromis* | Benthopelagic | Reef-associated | Planktivorous | Marine | ✓ | ✓ | ✓ | ✓ | ✓ | ✓ | ✓ | ✓ | ✓ | ✓ |
| *Clupanodon* | Pelagic | Pelagic | Planktivorous | Marine/Brackish/Freshwater | | | | | | ✓ | ✓ | ✓ | | ✓ |
| *Crenimugil* | Benthopelagic | Reef-associated | Omnivorous | Marine/Brackish/Freshwater | | | | | ✓ | ✓ | ✓ | ✓ | ✓ | ✓ |
| *Cryptocentrus* | Benthic | Reef-associated | Omnivorous | Marine | | ✓ | ✓ | ✓ | ✓ | ✓ | ✓ | | | ✓ |
| *Dascyllus* | Benthopelagic | Reef-associated | Omnivorous | Marine | | | | ✓ | | | | | | |
| *Decapterus* | Pelagic | Reef-associated | Invertivorous | Marine | | | | | | ✓ | ✓ | ✓ | ✓ | ✓ |
| *Dendrochirus* | Benthopelagic | Reef-associated | Invertivorous | Marine | | | ✓ | | | | | | | |
| *Deveximentum* | Pelagic | Reef-associated | Invertivorous | Marine/Brackish | | | | | | ✓ | ✓ | ✓ | ✓ | ✓ |
| *Diagramma* | Benthopelagic | Reef-associated | Invertivorous.Piscivorous | Marine | ✓ | ✓ | ✓ | ✓ | ✓ | ✓ | ✓ | ✓ | | ✓ |
| *Diploprion* | Benthopelagic | Reef-associated | Invertivorous.Piscivorous | Marine | | ✓ | | | ✓ | | | | | |
| *Drepane* | Benthopelagic | Reef-associated | Invertivorous.Piscivorous | Marine/Brackish/Freshwater | | | | | | | ✓ | ✓ | ✓ | ✓ |
| *Eleutheronema* | Benthopelagic | Demersal | Invertivorous.Piscivorous | Marine/Brackish/Freshwater | | | | | | ✓ | ✓ | ✓ | | ✓ |
| *Elops* | Benthopelagic | Pelagic | Invertivorous.Piscivorous | Marine/Brackish | | | | | | | ✓ | ✓ | ✓ | |
| *Encrasicholina* | Pelagic | Reef-associated | Invertivorous.Planktivorous | Marine/Brackish | | | | | | ✓ | ✓ | ✓ | ✓ | ✓ |
| *Enneapterygius* | Benthic | Reef-associated | Herbivorous | Marine | ✓ | | | | | ✓ | ✓ | ✓ | ✓ | ✓ |
| *Entomacrodus* | Benthic | Reef-associated | Herbivorous | Marine | | | | | | ✓ | ✓ | ✓ | ✓ | ✓ |
| *Epinephelus* | Benthopelagic | Reef-associated | Invertivorous.Piscivorous | Marine | | ✓ | | ✓ | ✓ | | ✓ | ✓ | | ✓ |
| *Equulites* | Benthopelagic | Demersal | Invertivorous | Marine | | | | | | ✓ | ✓ | ✓ | ✓ | ✓ |
| *Euthynnus* | Pelagic | Pelagic | Invertivorous.Piscivorous | Marine | | | | | | ✓ | ✓ | ✓ | ✓ | ✓ |
| *Eviota* | Benthic | Reef-associated | Omnivorous | Marine | | | | | ✓ | | | | | |
| *Fistularia* | Benthopelagic | Reef-associated | Invertivorous.Piscivorous | Marine | | ✓ | ✓ | | ✓ | | | | | |

| | | | | | | | | | | | | | | |
|---|---|---|---|---|---|---|---|---|---|---|---|---|---|---|
| *Gazza* | Benthopelagic | Reef-associated | Invertivorous | Marine/Brackish | | | | | | | | | ✓ | |
| *Gerres* | Benthopelagic | Reef-associated/Demersal | Invertivorous | Marine/Brackish/Freshwater | | ✓ | ✓ | ✓ | ✓ | ✓ | ✓ | ✓ | ✓ | ✓ |
| *Gymnothorax* | Benthic | Reef-associated | Invertivorous.Piscivorous | Marine | | ✓ | ✓ | | | ✓ | ✓ | ✓ | ✓ | |
| *Halichoeres* | Benthopelagic | Reef-associated | Invertivorous | Marine | ✓ | ✓ | ✓ | ✓ | ✓ | | | | | |
| *Hilsa* | Pelagic | Pelagic | Omnivorous | Marine/Brackish/Freshwater | | | | | | ✓ | ✓ | ✓ | ✓ | ✓ |
| *Hypoatherina* | Pelagic | Pelagic | Planktivorous | Marine | | | | | | ✓ | ✓ | ✓ | ✓ | ✓ |
| *Hyporhamphus* | Pelagic | Pelagic | Omnivorous | Marine/Brackish/Freshwater | | | | | | ✓ | ✓ | ✓ | ✓ | ✓ |
| *Istigobius* | Benthic | Reef-associated | Omnivorous | Marine | ✓ | ✓ | ✓ | ✓ | ✓ | ✓ | ✓ | ✓ | ✓ | ✓ |
| *Konosirus* | Pelagic | Pelagic | Planktivorous | Marine/Brackish | | | | | | ✓ | ✓ | ✓ | ✓ | ✓ |
| *Lagocephalus* | Benthopelagic | Demersal | Invertivorous | Marine/Brackish | | | | | | ✓ | ✓ | ✓ | ✓ | ✓ |
| *Larimichthys* | Benthopelagic | Demersal | Invertivorous.Piscivorous | Marine/Brackish | | | | | | ✓ | ✓ | ✓ | | ✓ |
| *Lepturacanthus* | Benthopelagic | Demersal | Invertivorous.Piscivorous | Marine/Brackish | | | | | | ✓ | ✓ | ✓ | ✓ | ✓ |
| *Lutjanus* | Benthopelagic | Reef-associated | Invertivorous.Piscivorous | Marine/Brackish | ✓ | ✓ | | ✓ | ✓ | ✓ | ✓ | ✓ | | ✓ |
| *Microcanthus* | Benthopelagic | Reef-associated | Omnivorous | Marine/Brackish | | ✓ | | ✓ | | | | | | |
| *Monacanthus* | Benthopelagic | Reef-associated | Omnivorous | Marine | | ✓ | ✓ | ✓ | ✓ | ✓ | ✓ | ✓ | ✓ | ✓ |
| *Mugil* | Benthopelagic | Demersal | Omnivorous | Marine/Brackish/Freshwater | | | | | | ✓ | ✓ | ✓ | ✓ | ✓ |
| *Myersina* | Benthic | Reef-associated | Omnivorous | Marine | | ✓ | | | | | | | | |
| *Nematalosa* | Pelagic | Demersal/Pelagic | Planktivorous | Marine/Brackish/Freshwater | | | | | | ✓ | ✓ | ✓ | ✓ | ✓ |
| *Neopomacentrus* | Benthopelagic | Reef-associated | Planktivorous | Marine | ✓ | ✓ | ✓ | ✓ | ✓ | ✓ | ✓ | ✓ | ✓ | ✓ |
| *Nibea* | Benthopelagic | Demersal | Invertivorous | Marine | | | | | | | ✓ | ✓ | ✓ | ✓ |
| *Nuchequula* | Pelagic | Pelagic | Invertivorous | Marine/Brackish | | | | | | ✓ | ✓ | ✓ | ✓ | ✓ |
| *Omobranchus* | Benthic | Reef-associated | Omnivorous | Marine/Brackish | ✓ | | | | | | | | | |
| *Ostorhinchus* | Benthopelagic | Reef-associated | Invertivorous.Piscivorous | Marine/Brackish | ✓ | ✓ | ✓ | ✓ | ✓ | ✓ | ✓ | ✓ | ✓ | ✓ |
| *Ostracion* | Benthopelagic | Reef-associated | Omnivorous | Marine | | | ✓ | | | | | | | |
| *Oxyurichthys* | Benthic | Demersal | Invertivorous | Marine/Brackish | | | | | | | | | ✓ | |
| *Pagrus* | Benthopelagic | Demersal | Invertivorous.Piscivorous | Marine | | ✓ | ✓ | | ✓ | | | | | |
| *Paracentropogon* | Benthic | Demersal | Invertivorous.Piscivorous | Marine | ✓ | | | | | | | | | |
| *Paramonacanthus* | Benthopelagic | Demersal | Invertivorous.Piscivorous | Marine | | ✓ | ✓ | | ✓ | | | | | |
| *Parapercis* | Benthic | Reef-associated | Invertivorous | Marine | | ✓ | ✓ | ✓ | ✓ | | | | | |
| *Parapristipoma* | Benthopelagic | Reef-associated | Planktivorous | Marine | | | | | | ✓ | ✓ | ✓ | ✓ | ✓ |
| *Parascorpaena* | Benthic | Reef-associated | Invertivorous | Marine/Brackish | | | | ✓ | | ✓ | ✓ | ✓ | | ✓ |
| *Parexocoetus* | Pelagic | Pelagic | Planktivorous | Marine | | | | | | ✓ | ✓ | ✓ | ✓ | ✓ |
| *Parioglossus* | Benthopelagic | Reef-associated | Planktivorous | Marine/Brackish | | | | | ✓ | | | | | |
| *Parupeneus* | Benthopelagic | Reef-associated | Invertivorous.Piscivorous | Marine/Brackish | | ✓ | ✓ | ✓ | ✓ | ✓ | ✓ | ✓ | ✓ | ✓ |
| *Pelates* | Benthopelagic | Reef-associated | Invertivorous.Piscivorous | Marine/Brackish | | | | | ✓ | | | | | |
| *Pempheris* | Benthopelagic | Reef-associated | Planktivorous | Marine | | ✓ | | ✓ | | ✓ | ✓ | ✓ | ✓ | ✓ |
| *Pennahia* | Benthopelagic | Demersal | Invertivorous.Piscivorous | Marine | | | | | | | | | ✓ | |
| *Petroscirtes* | Benthopelagic | Reef-associated | Omnivorous | Marine/Brackish | | ✓ | ✓ | | ✓ | ✓ | ✓ | | | ✓ |
| *Photopectoralis* | Benthopelagic | Demersal | Omnivorous | Marine/Brackish | | | | | | ✓ | ✓ | ✓ | ✓ | ✓ |
| *Planiliza* | Benthopelagic | Demersal | Omnivorous | Marine/Brackish/Freshwater | | | | ✓ | | ✓ | ✓ | ✓ | ✓ | ✓ |
| *Plectorhinchus* | Benthopelagic | Reef-associated | Invertivorous.Piscivorous | Marine/Brackish/Freshwater | | | | | | ✓ | | ✓ | | ✓ |
| *Plicomugil* | Benthopelagic | Reef-associated | Omnivorous | Marine | | | | | | ✓ | ✓ | ✓ | ✓ | ✓ |
| *Plotosus* | Benthopelagic | Reef-associated | Omnivorous | Marine/Brackish | ✓ | | | | | ✓ | ✓ | ✓ | | ✓ |
| *Pomacentrus* | Benthopelagic | Reef-associated | Herbivorous | Marine | | | | | ✓ | | | | | |
| *Pteragogus* | Benthopelagic | Reef-associated | Invertivorous | Marine | | | ✓ | | | | | | | |
| *Ptereleotris* | Benthopelagic | Reef-associated | Planktivorous | Marine | | | | | ✓ | | | | | |
| *Rastrelliger* | Pelagic | Pelagic | Planktivorous | Marine | | | | | | ✓ | ✓ | ✓ | ✓ | ✓ |

| | | | | | | | | | | | | | | |
|---|---|---|---|---|---|---|---|---|---|---|---|---|---|---|
| *Rhabdosargus* | Benthopelagic | Reef-associated | Omnivorous | Marine/Brackish | | | | | ✓ | ✓ | ✓ | ✓ | ✓ | ✓ |
| *Sardinella* | Pelagic | Pelagic | Planktivorous | Marine/Brackish | | | | | | ✓ | ✓ | ✓ | ✓ | ✓ |
| *Scarus* | Benthopelagic | Reef-associated | Herbivorous | Marine/Brackish | | | | ✓ | ✓ | | | | | |
| *Scatophagus* | Benthopelagic | Demersal | Omnivorous | Marine/Brackish/Freshwater | | | | | ✓ | | | | | |
| *Sciaenops* | Benthopelagic | Demersal | Invertivorous | Marine/Brackish | | | | | | ✓ | | ✓ | | ✓ |
| *Scorpaenopsis* | Benthic | Reef-associated | Invertivorous.Piscivorous | Marine | ✓ | | ✓ | ✓ | ✓ | | | | | |
| *Sebastiscus* | Benthic | Reef-associated | Invertivorous.Piscivorous | Marine | ✓ | ✓ | ✓ | | ✓ | ✓ | ✓ | ✓ | ✓ | ✓ |
| *Selar* | Pelagic | Reef-associated | Planktivorous | Marine | | | | | | | ✓ | ✓ | ✓ | ✓ |
| *Selaroides* | Benthopelagic | Reef-associated | Invertivorous.Piscivorous | Marine/Brackish | | ✓ | | ✓ | ✓ | | ✓ | ✓ | ✓ | ✓ |
| *Siganus* | Benthopelagic | Reef-associated | Herbivorous | Marine/Brackish | ✓ | ✓ | ✓ | ✓ | ✓ | ✓ | ✓ | ✓ | ✓ | ✓ |
| *Sillago* | Benthopelagic | Demersal | Invertivorous | Marine | ✓ | | | | | ✓ | ✓ | ✓ | ✓ | ✓ |
| *Sphyraena* | Benthopelagic | Reef-associated | Invertivorous.Piscivorous | Marine/Brackish | | | | | | ✓ | ✓ | ✓ | ✓ | ✓ |
| *Spratelloides* | Pelagic | Pelagic | Planktivorous | Marine | | | | | | ✓ | ✓ | ✓ | ✓ | ✓ |
| *Stethojulis* | Benthopelagic | Reef-associated | Invertivorous | Marine | | ✓ | ✓ | ✓ | ✓ | ✓ | ✓ | ✓ | ✓ | ✓ |
| *Stolephorus* | Pelagic | Pelagic | Planktivorous | Marine/Brackish | | | | | | ✓ | | ✓ | ✓ | ✓ |
| *Synodus* | Benthic | Reef-associated | Invertivorous.Piscivorous | Marine | | | ✓ | | ✓ | | | | | |
| *Taeniamia* | Benthopelagic | Reef-associated | Invertivorous | Marine | | ✓ | ✓ | ✓ | ✓ | ✓ | ✓ | ✓ | ✓ | ✓ |
| *Takifugu* | Benthopelagic | Reef-associated | Invertivorous | Marine | ✓ | ✓ | ✓ | ✓ | ✓ | ✓ | ✓ | ✓ | ✓ | ✓ |
| *Thalassoma* | Benthopelagic | Reef-associated | Invertivorous | Marine | | ✓ | ✓ | ✓ | ✓ | ✓ | ✓ | ✓ | ✓ | ✓ |
| *Thryssa* | Pelagic | Pelagic | Planktivorous | Marine/Brackish | | | | | | ✓ | ✓ | ✓ | ✓ | ✓ |
| *Thysanophrys* | Benthic | Reef-associated | Invertivorous.Piscivorous | Marine/Brackish | | | ✓ | ✓ | | | | | | |
| *Trachinotus* | Benthopelagic | Reef-associated | Invertivorous | Marine/Brackish | | | | | | ✓ | ✓ | ✓ | | ✓ |
| *Trichonotus* | Benthic | Reef-associated | Planktivorous | Marine/Brackish | | ✓ | | | | | | | | |
| *Tridentiger* | Benthic | Demersal | Invertivorous | Marine/Brackish/Freshwater | ✓ | | | | | ✓ | | ✓ | | ✓ |
| *Tylosurus* | Benthic | Reef-associated | Invertivorous.Piscivorous | Marine/Brackish | | | | | | ✓ | ✓ | ✓ | ✓ | ✓ |
| *Upeneus* | Benthopelagic | Reef-associated/Demersal | Invertivorous | Marine/Brackish | | ✓ | ✓ | ✓ | ✓ | ✓ | ✓ | ✓ | ✓ | ✓ |
| *Valenciennea* | Benthopelagic | Reef-associated | Invertivorous | Marine | | | | ✓ | ✓ | | | | | |
| *Zanclus* | Benthopelagic | Reef-associated | Invertivorous | Marine | | | ✓ | | | | | | | |

Supplementary Table 4. Global and regional differences in the (a) total number and (b) average and standard deviation (Std.) of fish genera recorded between regions across both and separate sampling seasons using UVC and eDNA metabarcoding.

(a)

| | *Total number* | | | | | |
|---|---|---|---|---|---|---|
| | UVC | | | eDNA metabarcoding | | |
| Region | Wet + Dry | Wet | Dry | Wet + Dry | Wet | Dry |
| Lantau | 19 | 15 | 10 | 75 | 60 | 61 |
| South Hong Kong (S-HK) | 37 | 34 | 16 | 77 | 58 | 63 |
| Southeast Hong Kong (SE-HK) | 38 | 33 | 18 | 77 | 61 | 61 |
| East Sai Kung (E-SK) | 37 | 35 | 20 | 65 | 51 | 39 |
| North Sai Kung (N-SK) | 52 | 49 | 27 | 78 | 56 | 66 |
| *Across regions* | *73* | *67* | *43* | *84* | *64* | *73* |

(b)

| | *Mean ± Std.* | | | | | |
|---|---|---|---|---|---|---|
| | UVC | | | eDNA metabarcoding | | |
| Region | Wet + Dry | Wet | Dry | Wet + Dry | Wet | Dry |
| Lantau | 3.8 ± 2.3 | 4.4 ± 2.3 | 3.2 ± 2.4 | 50.9 ± 5.7 | 47.6 ± 6.4 | 54.2 ± 2.2 |
| South Hong Kong (S-HK) | 12.0 ± 6.2 | 17.4 ± 3.2 | 6.6 ± 1.9 | 50.9 ± 5.4 | 46.4 ± 3.6 | 55.4 ± 1.1 |
| Southeast Hong Kong (SE-HK) | 10.2 ± 5.7 | 14.4 ± 4.6 | 6.0 ± 2.5 | 51.4 ± 2.6 | 50.6 ± 3.0 | 52.2 ± 2.2 |
| East Sai Kung (E-SK) | 11.9 ± 4.5 | 15.2 ± 2.9 | 8.6 ± 3.0 | 29.5 ± 7.7 | 35.2 ± 4.1 | 23.8 ± 5.8 |
| North Sai Kung (N-SK) | 16.4 ± 6.9 | 22.6 ± 1.8 | 10.2 ± 2.6 | 46.9 ± 8.2 | 40.8 ± 4.9 | 53.0 ± 5.7 |
| *Across regions* | *10.9 ± 6.6* | *14.8 ± 6.7* | *6.9 ± 3.3* | *45.9 ± 10.3* | *44.1 ± 7.0* | *47.7 ± 12.8* |

Supplementary Table 5. PERMANOVA and PERMDISP tests based on 9999 permutations for pairwise comparisons of fish composition among regions for both seasons and during either season for (a) underwater visual census (UVC) and (b) eDNA metabarcoding. *df*: degree of freedom; Significant results are in bold. ***: $p < 0.001$; **: $p < 0.01$; *: $p < 0.05$.

*a) UVC*

| **Dataset** | **Group** | **PERMANOVA** | | | | | **PERMDISP** | | | | |
|---|---|---|---|---|---|---|---|---|---|---|---|
| | | ***df*** | $R^2$ | ***Pseudo-F statistic*** | ***p value*** | | $df_1$ | $df_2$ | ***F statistic*** | ***p value*** | |
| Wet + Dry | | | | | | | | | | | |
| | Region | 4 | 0.21 | 3.185 | **< 0.001** | *** | 4 | 45 | 4.055 | **0.007** | ** |
| | Season | 1 | 0.05 | 2.894 | **< 0.001** | *** | 1 | 48 | 4.040 | **0.049** | * |
| | Region x Season | 4 | 0.09 | 1.313 | **0.032** | * | | | | | |
| | Residual | 40 | 0.65 | | | | | | | | |
| | Total | 49 | 1.00 | | | | | | | | |
| Wet | | | | | | | | | | | |
| | Region | 4 | 0.32 | 2.402 | **< 0.001** | *** | 4 | 20 | 4.203 | **0.012** | * |
| | Residual | 20 | 0.68 | | | | | | | | |
| | Total | 24 | 1.00 | | | | | | | | |
| Dry | | | | | | | | | | | |
| | Region | 4 | 0.30 | 2.127 | **< 0.001** | *** | 4 | 20 | 0.877 | 0.483 | |
| | Residual | 20 | 0.70 | | | | | | | | |
| | Total | 24 | 1.00 | | | | | | | | |

*b) eDNA metabarcoding*

| **Dataset** | **Group** | **PERMANOVA** | | | | | **PERMDISP** | | | | |
|---|---|---|---|---|---|---|---|---|---|---|---|
| | | ***df*** | $R^2$ | ***Pseudo-F statistic*** | ***p value*** | | $df_1$ | $df_2$ | ***F statistic*** | ***p value*** | |
| Wet + Dry | | | | | | | | | | | |
| | Region | 4 | 0.31 | 15.106 | **< 0.001** | *** | 4 | 45 | 6.067 | **0.003** | ** |
| | Season | 1 | 0.30 | 59.919 | **< 0.001** | *** | 1 | 48 | 0.091 | 0.768 | |
| | Region x Season | 4 | 0.19 | 9.456 | **< 0.001** | *** | | | | | |
| | Residual | 40 | 0.20 | | | | | | | | |
| | Total | 49 | 1.00 | | | | | | | | |
| Wet | | | | | | | | | | | |
| | Region | 4 | 0.64 | 8.754 | **< 0.001** | *** | 4 | 20 | 3.276 | **0.028** | * |
| | Residual | 20 | 0.36 | | | | | | | | |
| | Total | 24 | 1.00 | | | | | | | | |
| Dry | | | | | | | | | | | |
| | Region | 4 | 0.80 | 19.868 | **< 0.001** | *** | 4 | 20 | 21.875 | **< 0.001** | *** |
| | Residual | 20 | 0.20 | | | | | | | | |
| | Total | 24 | 1.00 | | | | | | | | |

Supplementary Table 6. Pairwise comparisons for PERMANOVA – *PairwiseAdonis* and for PERMDISP – *Betadisper* based on 9999 permutations among regions for both sampling seasons combined and for only wet and dry seasons for (a-c) underwater visual census (UVC) and (d-f) eDNA metabarcoding. S-HK: South Hong Kong, SE-HK: Southeast Hong Kong, E-SK: East Sai Kung, N-SK: North Sai Kung. Significant results are presented as ***: $p < 0.001$; **: $p < 0.01$; *: $p < 0.05$.

*a) Wet + Dry (UVC)*

| | | ***Betadisper*** | | | | |
|---|---|---|---|---|---|---|
| | ***Region*** | Lantau | S-HK | SE-HK | E-SK | N-SK |
| ***PairwiseAdonis*** | Lantau | / | * | | ** | ** |
| | S-HK | ** | / | | | |
| | SE-HK | *** | | / | | |
| | E-SK | *** | | * | / | |
| | N-SK | *** | ** | ** | * | / |

*b) Wet (UVC)*

| | | ***Betadisper*** | | | | |
|---|---|---|---|---|---|---|
| | ***Region*** | Lantau | S-HK | SE-HK | E-SK | N-SK |
| ***PairwiseAdonis*** | Lantau | / | ** | | ** | ** |
| | S-HK | | / | | | |
| | SE-HK | | | / | | |
| | E-SK | | | | / | |
| | N-SK | | | | | / |

*c) Dryr (UVC)*

| | | ***Betadisper*** | | | | |
|---|---|---|---|---|---|---|
| | ***Region*** | Lantau | S-HK | SE-HK | E-SK | N-SK |
| ***PairwiseAdonis*** | Lantau | / | | | | |
| | S-HK | | / | | | |
| | SE-HK | | | / | | |
| | E-SK | | | | / | |
| | N-SK | | | | | / |

*d) Wet + Dry (eDNA metabarcoding)*

***Betadisper*** (columns); ***PairwiseAdonis*** (rows)

| ***Region*** | Lantau | S-HK | SE-HK | E-SK | N-SK |
|---|---|---|---|---|---|
| Lantau | / | | | ** | |
| S-HK | | / | | ** | |
| SE-HK | | | / | *** | |
| E-SK | *** | ** | *** | / | * |
| N-SK | | | | *** | / |

*e) Wet (eDNA metabarcoding)*

***Betadisper*** (columns); ***PairwiseAdonis*** (rows)

| ***Region*** | Lantau | S-HK | SE-HK | E-SK | N-SK |
|---|---|---|---|---|---|
| Lantau | / | | | | |
| S-HK | | / | | | ** |
| SE-HK | | | / | | ** |
| E-SK | | | | / | |
| N-SK | | | | | / |

*f) Dry (eDNA metabarcoding)*

***Betadisper*** (columns); ***PairwiseAdonis*** (rows)

| ***Region*** | Lantau | S-HK | SE-HK | E-SK | N-SK |
|---|---|---|---|---|---|
| Lantau | / | | | *** | |
| S-HK | | / | | *** | |
| SE-HK | | | / | *** | |
| E-SK | | | | / | *** |
| N-SK | | | | | / |

Supplementary Table 7. Indicator species analysis (ISA) results for the combinations of region and sampling seasons based on underwater visual census data. Significant results are presented as ***: $p < 0.001$; **: $p < 0.01$; *: $p < 0.05$.

```
Multilevel pattern analysis
 ---------------------------

 Association function: r.g
 Significance level (alpha): 0.05

 Total number of species: 73
 Selected number of species: 21
 Number of species associated to 1 group: 7
 Number of species associated to 2 groups: 2
 Number of species associated to 3 groups: 1
 Number of species associated to 4 groups: 4
 Number of species associated to 5 groups: 2
 Number of species associated to 6 groups: 3
 Number of species associated to 7 groups: 1
 Number of species associated to 8 groups: 1
 Number of species associated to 9 groups: 0

 List of species associated to each combination:

 Group Wet.S-HK  #sps.  1
           stat p.value
Diploprion 0.639   0.021 *

 Group Wet.E-SK  #sps.  1
            stat p.value
Monacanthus 0.538    0.05 *

 Group Wet.N-SK  #sps.  5
             stat p.value
Parioglossus 0.758   0.008 **
Ptereleotris 0.758   0.003 **
Rhabdosargus 0.758   0.003 **
Petroscirtes 0.634   0.008 **
Scarus       0.556   0.040 *

 Group Dry.N-SK+Wet.N-SK  #sps.  2
              stat p.value
Eviota        0.667   0.002 **
Asterropteryx 0.585   0.025 *

 Group Wet.S-HK+Wet.SE-HK+Wet.N-SK  #sps.  1
                   stat p.value
Apogonichthyoides 0.566    0.01 **

 Group Dry.S-HK+Wet.S-HK+Wet.SE-HK+Wet.E-SK  #sps.  1
        stat p.value
Chromis 0.653   0.001 ***
```

```
 Group Wet.S-HK+Wet.SE-HK+Wet.E-SK+Wet.N-SK  #sps.  1
              stat p.value
Cephalopholis 0.624   0.005 **

 Group Wet.S-HK+Dry.E-SK+Wet.E-SK+Wet.N-SK  #sps.  1
            stat p.value
Stethojulis 0.592   0.005 **

 Group Dry.E-SK+Wet.E-SK+Dry.N-SK+Wet.N-SK  #sps.  1
            stat p.value
Amblygobius 0.84   0.001 ***

 Group Wet.S-HK+Wet.SE-HK+Dry.E-SK+Wet.E-SK+Wet.N-SK  #sps.  1
           stat p.value
Parupeneus 0.689   0.001 ***

 Group Wet.S-HK+Wet.SE-HK+Wet.E-SK+Dry.N-SK+Wet.N-SK  #sps.  1
           stat p.value
Parapercis 0.667   0.002 **

 Group Dry.S-HK+Wet.S-HK+Wet.SE-HK+Wet.E-SK+Dry.N-SK+Wet.N-SK  #sps.  1
                stat p.value
Neopomacentrus 0.707   0.001 ***

 Group Wet.S-HK+Wet.SE-HK+Dry.E-SK+Wet.E-SK+Dry.N-SK+Wet.N-SK  #sps.  2
          stat p.value
Abudefduf 0.612   0.001 ***
Gerres    0.612   0.006 **

 Group Dry.S-HK+Wet.S-HK+Dry.SE-HK+Wet.SE-HK+Dry.E-SK+Wet.E-SK+Wet.N-SK  #sps.  1
           stat p.value
Amphiprion 0.535   0.015 *

 Group Dry.S-HK+Wet.S-HK+Dry.SE-HK+Wet.SE-HK+Dry.E-SK+Wet.E-SK+Dry.N-SK+Wet.N-SK  #sps.  1
            stat p.value
Halichoeres 0.655   0.003 **
---
Signif. codes:  0 ‘***’ 0.001 ‘**’ 0.01 ‘*’ 0.05 ‘.’ 0.1 ‘ ’ 1
```

Supplementary Table 8. Indicator species analysis (ISA) results for the combinations of region and sampling seasons based on eDNA metabarcoding data. Significant results are presented as ***: $p < 0.001$; **: $p < 0.01$; *: $p < 0.05$.

```
Multilevel pattern analysis
 ---------------------------

 Association function: r.g
 Significance level (alpha): 0.05

 Total number of species: 79
 Selected number of species: 76
 Number of species associated to 1 group: 2
 Number of species associated to 2 groups: 10
 Number of species associated to 3 groups: 5
 Number of species associated to 4 groups: 15
 Number of species associated to 5 groups: 12
 Number of species associated to 6 groups: 12
 Number of species associated to 7 groups: 5
 Number of species associated to 8 groups: 6
 Number of species associated to 9 groups: 9

 List of species associated to each combination:

 Group Dry.E-SK  #sps.  1
       stat p.value
Gazza 0.758   0.005 **

 Group Wet.E-SK  #sps.  1
       stat p.value
Elops 0.826   0.001 ***

 Group Dry.Lantau+Dry.SE-HK  #sps.  3
                stat p.value
Plectorhinchus 0.941   0.001 ***
Sciaenops      0.941   0.001 ***
Larimichthys   0.890   0.001 ***

 Group Wet.Lantau+Wet.SE-HK  #sps.  3
             stat p.value
Alectis     0.890   0.001 ***
Chaetodon   0.844   0.001 ***
Gymnothorax 0.802   0.001 ***

 Group Dry.S-HK+Dry.N-SK  #sps.  4
              stat p.value
Cryptocentrus 0.941   0.001 ***
Petroscirtes  0.941   0.001 ***
Epinephelus   0.890   0.001 ***
Nibea         0.821   0.001 ***
```

Group Wet.Lantau+Wet.SE-HK+Wet.N-SK #sps. 1
stat p.value
Tridentiger 0.764 0.001 ***

Group Dry.S-HK+Dry.E-SK+Dry.N-SK #sps. 1
stat p.value
Drepane 0.706 0.002 **

Group Dry.S-HK+Wet.E-SK+Dry.N-SK #sps. 1
stat p.value
Selaroides 0.767 0.001 ***

Group Wet.S-HK+Wet.SE-HK+Wet.N-SK #sps. 1
stat p.value
Selar 0.706 0.001 ***

Group Wet.S-HK+Dry.E-SK+Wet.N-SK #sps. 1
stat p.value
Thalassoma 0.81 0.001 ***

Group Dry.Lantau+Wet.Lantau+Dry.SE-HK+Wet.SE-HK #sps. 1
stat p.value
Lagocephalus 0.757 0.001 ***

Group Dry.Lantau+Dry.S-HK+Dry.SE-HK+Dry.E-SK #sps. 1
stat p.value
Chromis 0.753 0.001 ***

Group Dry.Lantau+Dry.S-HK+Dry.SE-HK+Wet.E-SK #sps. 1
stat p.value
Photopectoralis 0.622 0.002 **

Group Dry.Lantau+Dry.S-HK+Dry.SE-HK+Dry.N-SK #sps. 9
stat p.value
Eleutheronema 1.000 0.001 ***
Lutjanus 1.000 0.001 ***
Parascorpaena 1.000 0.001 ***
Parapristipoma 0.959 0.001 ***
Plotosus 0.959 0.001 ***
Acanthopagrus 0.919 0.001 ***
Clupanodon 0.919 0.001 ***
Lepturacanthus 0.793 0.001 ***
Stethojulis 0.667 0.002 **

Group Wet.Lantau+Dry.S-HK+Wet.SE-HK+Dry.N-SK #sps. 1
stat p.value
Amphiprion 0.794 0.001 ***

Group Wet.Lantau+Wet.S-HK+Wet.SE-HK+Wet.E-SK #sps. 2
stat p.value
Parexocoetus 0.875 0.001 ***
Carangoides 0.791 0.001 ***

Group Dry.Lantau+Wet.Lantau+Wet.S-HK+Wet.SE-HK+Wet.N-SK #sps. 1
stat p.value

Ambassis 0.682   0.001 ***

Group Dry.Lantau+Wet.Lantau+Dry.SE-HK+Wet.SE-HK+Dry.N-SK  #sps.  1
stat p.value
Nuchequula 0.725   0.001 ***

Group Dry.Lantau+Wet.Lantau+Dry.SE-HK+Wet.SE-HK+Wet.N-SK  #sps.  1
stat p.value
Stolephorus 0.843   0.001 ***

Group Dry.Lantau+Dry.S-HK+Dry.SE-HK+Wet.SE-HK+Dry.N-SK  #sps.  1
stat p.value
Bathygobius 0.633   0.002 **

Group Dry.Lantau+Wet.S-HK+Dry.SE-HK+Wet.E-SK+Wet.N-SK  #sps.  1
stat p.value
Monacanthus 0.667   0.001 ***

Group Wet.Lantau+Dry.S-HK+Wet.S-HK+Dry.N-SK+Wet.N-SK  #sps.  1
stat p.value
Asterropteryx 0.881   0.001 ***

Group Wet.Lantau+Wet.S-HK+Wet.SE-HK+Wet.E-SK+Wet.N-SK  #sps.  4
stat p.value
Euthynnus    0.886   0.001 ***
Hyporhamphus 0.886   0.001 ***
Taeniamia    0.783   0.001 ***
Tylosurus    0.750   0.001 ***

Group Dry.S-HK+Wet.S-HK+Dry.E-SK+Dry.N-SK+Wet.N-SK  #sps.  2
stat p.value
Amblygobius 0.801   0.001 ***
Sphyraena   0.700   0.001 ***

Group Dry.Lantau+Wet.Lantau+Dry.S-HK+Dry.SE-HK+Wet.SE-HK+Dry.N-SK  #sps.  5
stat p.value
Trachinotus 0.885   0.001 ***
Callionymus 0.839   0.001 ***
Upeneus     0.834   0.001 ***
Diagramma   0.803   0.001 ***
Sillago     0.633   0.004 **

Group Dry.Lantau+Wet.Lantau+Wet.S-HK+Dry.SE-HK+Wet.SE-HK+Wet.N-SK  #sps.  1
stat p.value
Konosirus 0.879   0.001 ***

Group Dry.Lantau+Dry.S-HK+Wet.S-HK+Dry.SE-HK+Wet.E-SK+Dry.N-SK  #sps.  1
stat p.value
Istigobius 0.624   0.004 **

Group Dry.Lantau+Dry.S-HK+Wet.S-HK+Dry.SE-HK+Dry.N-SK+Wet.N-SK  #sps.  1
stat p.value
Arothron 0.839   0.001 ***

Group Dry.Lantau+Dry.S-HK+Dry.SE-HK+Dry.E-SK+Wet.E-SK+Dry.N-SK  #sps.  1
stat p.value

Equulites 0.713   0.001 ***

Group Dry.Lantau+Wet.S-HK+Dry.SE-HK+Wet.SE-HK+Wet.E-SK+Wet.N-SK  #sps.  1
stat p.value
Rhabdosargus 0.582   0.003 **

Group Wet.Lantau+Wet.S-HK+Wet.SE-HK+Dry.E-SK+Wet.E-SK+Wet.N-SK  #sps.  2
stat p.value
Rastrelliger  0.885   0.001 ***
Spratelloides 0.764   0.001 ***

Group Dry.Lantau+Wet.Lantau+Dry.S-HK+Dry.SE-HK+Wet.SE-HK+Dry.E-SK+Dry.N-SK  #sps.  1
stat p.value
Deveximentum 0.764   0.001 ***

Group Dry.Lantau+Wet.Lantau+Wet.S-HK+Dry.SE-HK+Wet.SE-HK+Wet.E-SK+Wet.N-SK  #sps.  1
stat p.value
Decapterus 0.782   0.001 ***

Group Dry.Lantau+Dry.S-HK+Wet.S-HK+Dry.SE-HK+Wet.E-SK+Dry.N-SK+Wet.N-SK  #sps.  1
stat p.value
Plicomugil 0.654   0.002 **

Group Wet.Lantau+Dry.S-HK+Wet.S-HK+Wet.SE-HK+Wet.E-SK+Dry.N-SK+Wet.N-SK  #sps.  1
stat p.value
Parupeneus 0.706   0.002 **

Group Dry.S-HK+Wet.S-HK+Wet.SE-HK+Dry.E-SK+Wet.E-SK+Dry.N-SK+Wet.N-SK  #sps.  1
stat p.value
Ostorhinchus 0.656   0.001 ***

Group Dry.Lantau+Wet.Lantau+Dry.S-HK+Wet.S-HK+Dry.SE-HK+Wet.SE-HK+Dry.N-SK+Wet.N-SK  #sps.  3
stat p.value
Planiliza    0.821   0.001 ***
Thryssa      0.807   0.002 **
Hypoatherina 0.690   0.002 **

Group Dry.Lantau+Wet.Lantau+Dry.S-HK+Wet.S-HK+Wet.SE-HK+Dry.E-SK+Wet.E-SK+Dry.N-SK  #sps.  1
stat p.value
Hilsa 0.59   0.047 *

Group Dry.Lantau+Wet.Lantau+Dry.S-HK+Dry.SE-HK+Wet.SE-HK+Dry.E-SK+Wet.E-SK+Dry.N-SK  #sps.  1
stat p.value
Crenimugil 0.807   0.001 ***

Group Wet.Lantau+Dry.S-HK+Wet.S-HK+Wet.SE-HK+Dry.E-SK+Wet.E-SK+Dry.N-SK+Wet.N-SK  #sps.  1
stat p.value
Alepes 0.807   0.001 ***

Group Dry.Lantau+Wet.Lantau+Dry.S-HK+Wet.S-HK+Dry.SE-HK+Wet.SE-HK+Dry.E-SK+Dry.N-SK+Wet.N-SK  #sps.  1
stat p.value
Mugil 0.885   0.002 **

Group Dry.Lantau+Wet.Lantau+Dry.S-HK+Wet.S-HK+Dry.SE-HK+Wet.SE-HK+Wet.E-SK+Dry.N-SK+Wet.N-SK  #sps.  8

| | stat | p.value | |
|---|---|---|---|
| Atherinomorus | 1.000 | 0.001 | *** |
| Cephalopholis | 1.000 | 0.001 | *** |
| Nematalosa | 0.885 | 0.001 | *** |
| Takifugu | 0.826 | 0.002 | ** |
| Enneapterygius | 0.778 | 0.002 | ** |
| Neopomacentrus | 0.764 | 0.002 | ** |
| Sebastiscus | 0.667 | 0.003 | ** |
| Entomacrodus | 0.639 | 0.021 | * |

---

Signif. codes: 0 ‘***’ 0.001 ‘**’ 0.01 ‘*’ 0.05 ‘.’ 0.1 ‘ ’ 1
